# Utilizing a nanobody recruitment approach for assessing serine palmitoyltransferase activity in ER sub-compartments of yeast

**DOI:** 10.1101/2023.03.29.534722

**Authors:** Bianca M. Esch, Stefan Walter, Oliver Schmidt, Florian Fröhlich

## Abstract

Sphingolipids (SP) are one of the three major lipid classes in eukaryotic cells and serve as structural components of the plasma membrane. The rate-limiting step in SP biosynthesis is catalyzed by serine palmitoyltransferase (SPT). In yeast, SPT consists of two catalytic subunits (Lcb1 and Lcb2), a regulatory subunit (Tsc3), negative regulators (Orm1 and Orm2), and the phosphatidylinositol-4-phosphate (PI4P) phosphatase Sac1, collectively known as the SPOTS complex. Regulating SPT activity enables cells to adapt SP metabolism to changing environmental conditions. Therefore, the Orm proteins are phosphorylated by two signaling pathways originating from either the plasma membrane localized target of rapamycin (TOR) complex 2 or the lysosomal/vacuolar TOR complex 1. Moreover, uptake of exogenous serine is necessary for the regulation of SP biosynthesis, which suggests the existence of differentially regulated SPT pools based on their intracellular localization. However, tools for measuring lipid metabolic enzyme activity in different cellular compartments are currently not available. We have developed a nanobody recruitment system that enables the re-localization of the SPOTS complex to the nuclear or peripheral ER. By combining this system with sphingolipid flux analysis, we have identified two distinct active SPT pools in cells. Our method thus serves as a new and versatile tool to measure lipid metabolism with sub-cellular resolution.

## Introduction

Sphingolipids (SPs) are an essential class of lipids mainly found in the outer leaflet of the plasma membrane. They can act as signaling molecules as well as structural components of the plasma membrane (van Meer *et al*, 2008; Cartier & Hla, 2019). In order to allow cells and membranes to adapt, SP levels respond to different environmental conditions. For example, SP levels in yeast are changed in different carbon sources (Klose *et al*, 2012) and exposure of cells to heat rapidly elevates SP levels (Dickson *et al*, 1997). In mammalian cells, changes in SP composition in response to external stimuli, for example the amount of carbon source (Mondal *et al*, 2022), are also known. In particular, under conditions where increased amounts of sphingolipids are required, as in the formation of the myelin sheath, increased SP biosynthesis is observed (Davis *et al*, 2020).

SP biosynthesis begins in the endoplasmic reticulum (ER) with the enzyme serine palmitoyltransferase (SPT) (Buede *et al*, 1991; Gable *et al*, 2000; Nagiec *et al*, 1994). SPT catalyzes the first and rate-limiting step of SP biosynthesis by condensing L-serine and palmitoyl-CoA to form 3-ketosphinganine (3-KS). This short-lived intermediate is reduced to dihydrosphingosine (DHS), which in yeast is further hydroxylated to form phytosphingosine (PHS). DHS and PHS together are also known as long chain bases (LCBs). Very long chain fatty acids (VLCFAs) with 24 or 26 carbon atoms are then amide-linked to LCBs to form ceramides (D’mello *et al*, 1994; Guillas, 2001; Vallée & Riezman, 2005). Ceramides are transported to the Golgi apparatus through vesicular and non-vesicular transport (Kajiwara *et al*, 2013; Liu *et al*, 2017; Ikeda *et al*, 2020; Limar *et al*, 2023). In the Golgi apparatus, inositol containing head groups are added to form complex sphingolipids, which are then transported to the plasma membrane (Klemm *et al*, 2009).

SP biosynthesis is highly regulated with multiple input signals. The SPT in yeast forms together with the Orm proteins Orm1 and Orm2, the small subunit Tsc3 and the phosphatidylinositol-4-phosphate (PI4P) phosphatase Sac1 the SPOTS complex (Breslow *et al*, 2010; Han *et al*, 2010). While the mammalian SPT acts as a dimer, we have recently postulated that the yeast complex forms a monomer with either Orm1 or Orm2 bound to it (Schäfer *et al*, 2023). In yeast, SPT activity is regulated by phosphorylation of the Orm proteins. Orm phosphorylation originates either from the plasma membrane localized target of rapamycin complex 2 (TORC2) signaling pathway via the Ypk kinases 1 and 2 or from the TORC1 signaling pathway starting at the vacuole (Berchtold *et al*, 2012; Tabuchi *et al*, 2006; Roelants *et al*, 2011; Niles & Powers, 2012; Shimobayashi *et al*, 2013). In addition, the SPT is regulated via the downstream metabolite ceramide itself, which is also dependent on the Orm proteins (Davis *et al*, 2019). The two Orm proteins also underlie different regulatory mechanisms. Only the Orm2 protein is a substrate for the endosome/Golgi associated degradation (EGAD) pathway that depends on its phosphorylation by Ypk1/2, export from the ER, ubiquitination and finally degradation via the proteasome (Schmidt *et al*, 2019; Bhaduri *et al*, 2023). In addition, it was suggested that upregulation of SPT activity is directly linked to the uptake of exogenous serine via the Gnp1 amino acid permease (Esch *et al*, 2020). Other regulatory mechanisms controlling sphingolipid biogenesis involve the regulation of VLCFA biosynthesis (Olson *et al*, 2015; Zimmermann *et al*, 2013) and the post-translational regulation of the ceramide synthase (Muir *et al*, 2014).

Together, this highlights a complex regulatory network of SP biosynthesis regulation. However, the molecular mechanism underlying phosphorylation mediated regulation of the Orm proteins still remains enigmatic. It is not clear how the rapid increase in LCB biosynthesis upon heat shock can be aligned with the relatively slow EGAD degradation pathway of Orm2. In addition to the temporal regulation, spatial components of SPT regulation were not studied at all.

To shed light on these processes, we developed a nanobody based tool allowing us to recruit the SPOTS complex to different sub-compartments of the ER, namely the nuclear and the peripheral ER. We use multiple controls to determine the functionality of the system. It allows us to measure protein-protein interactions *in vivo*. By combining this re-localization tool with SP biosynthesis flux analysis, we are able to determine differentially active pools of the SPT in sub-cellular compartments. We believe that our study will be a starting point for the analysis of lipid metabolism with sub-cellular resolution.

## Results

Despite multiple efforts, the exact mechanism of Orm-dependent regulation of SPT remains enigmatic. Most experiments regarding Orm protein phosphorylation and its effect on SPT activity have been conducted in *ORM* deletion strains or in the presence of sphingolipid metabolism inhibitors, such as myriocin. However, inhibiting the enzyme whose Orm-dependent regulation should be investigated poses problems in itself. In addition to the complicated regulatory mechanism, we previously proposed that spatially separated SPT pools, specifically the nuclear envelope resident pool and the peripheral ER resident pool, are differentially regulated (Esch *et al*, 2020). We now aimed to study sphingolipid biosynthesis regulation under heat shock, a physiological condition known to rapidly upregulate serine palmitoyltransferase activity, with sub-cellular resolution.

First, we compared the levels of LCBs under control conditions and after a 5-minute heat shock in WT cells using targeted lipidomics. As previously reported (Dickson *et al*, 1997), we measured a rapid four-fold increase in LCBs after the short treatment **(Fig. 1a)**. We had previously demonstrated that increased *de novo* LCB biosynthesis is directly dependent on the uptake of exogenous serine through the general amino acid permease Gnp1. Consistent with our previous findings, deletion of *GNP1* resulted in a blunted heat shock response, while deletion of the endogenous serine biosynthesis pathway (*ser2Δ*) had no effect on LCB biosynthesis **(Fig. 1a)**. The incomplete block of heat shock induced LCB biosynthesis in a *gnp1Δ* strain may be due to the presence of the Gnp1 homolog Agp1. In summary, our experiments demonstrate that a short heat shock enables the investigation of SPT regulation in yeast cells.

**Figure 1:**
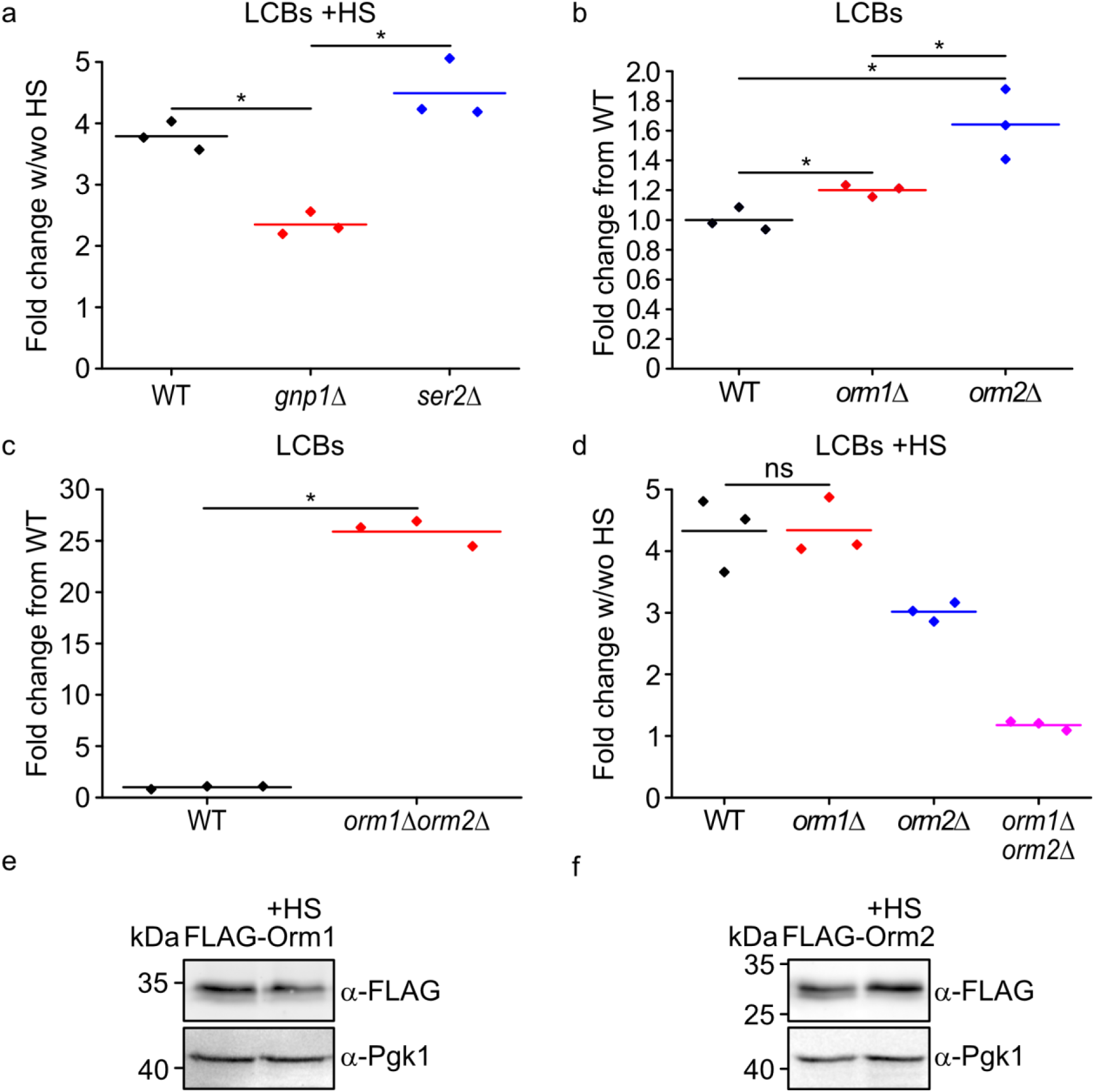
Orm proteins mediate SPT upregulation after heat shock. **(a)** Serine uptake by Gnp1 facilitates SPT upregulation. Mass spectrometry-based analysis of LCBs with and without 5 minutes of heat shock at 39° C. Displayed are the fold changes of LCBs for cells with versus without heat shock treatment in WT (black), gnp1Δ (red) and ser2Δ (blue) cells. Dots correspond to the values of three independent experiments. *, P-value <0.05, calculated from t-test. **(b)** One Orm protein can substitute for the corresponding other Orm protein. Mass spectrometry-based analysis of LCBs. Displayed are the fold changes from WT of WT (black), orm1Δ (red) and orm2Δ (blue) cells. Dots correspond to the values of three independent experiments. *, P-value <0.05, calculated from t-test. **(c)** LCBs levels are highly increased after ORM1 and ORM2 deletion. Mass spectrometry-based analysis of LCBs. Displayed are the fold changes from WT of WT (black), and orm1Δorm2Δ (red) cells. Dots correspond to the values of three independent experiments. *, P-value <0.05, calculated from t-test. **(d)** Orm proteins mediate LCB upregulation after heat shock. Mass spectrometry-based analysis of LCBs with and without 5 minutes of heat shock at 39° C. Displayed are the fold changes of LCBs for cells with versus without heat shock treatment in WT (black), orm1Δ (red), orm2Δ (blue) and orm1Δorm2Δ (pink) cells. Dots correspond to the values of three independent experiments. *, P-value <0.05, calculated from t-test. **(e, f)** Orm2 phosphorylation increases after heat shock. Phosphorylation pattern of 3xFLAG-Orm1 **(e)** and 3xFLAG-Orm2 **(f)** with and without heat shock. Cells were grown in YPD and were subjected to heat shock for 5 minutes at 39°C (+HS) or kept at room temperature. Equal amounts of cells were lysed and analyzed by western blotting using antibodies against the FLAG-tag or Pgk1 as a loading control.

Next, we assessed the individual contribution of the Orm proteins to heat shock dependent LCB up-regulation. Deletion of *ORM2* resulted in a significant increase in steady-state LCB levels compared to WT control cells. Consistent with previous findings, we detected a small but significant increase in LCB levels after deletion of *ORM1,* which was less pronounced than the effect of *orm2Δ* (Han *et al*, 2010) **(Fig. 1b)**. In contrast, deletion of both *ORM* genes led to a 25-fold increase in LCBs as previously reported (Breslow *et al*, 2010) (**Fig. 1c**). Taken together, these results suggest that each Orm1 or Orm2 protein is sufficient to maintain LCB levels within a physiological range under basal conditions. To investigate whether each Orm protein is also capable of maintaining LCB levels under rapidly changing conditions, we exposed *orm1Δ*, *orm2Δ* and *orm1Δorm2Δ* cells to heat shock, measured their LCB levels, and compared their heat shock mediated increase of LCBs to those of a WT yeast strain **(Fig. 1d)**. Both single deletion strains were still able to respond to the temperature change with a rapid increase in LCB biosynthesis. However, heat shock dependent LCB increase was significantly lower in *orm2*Δ cells with a 3-fold increase compared to a 4-fold increase in *orm1*Δ and WT cells. In contrast, the *orm1Δorm2Δ* strain did not show any further increase in LCB levels after heat shock **(Fig. 1d)**. Taken together, this suggests that both Orm proteins contribute to increased LCB biosynthesis after heat shock, with Orm2 making the greater contribution.

Subsequently, we evaluated whether the short 5-minute heat shock resulted in increased phosphorylation of the negative SPT regulators Orm1 and Orm2. We observed increased phosphorylation for Orm2 but not for Orm1 after heat shock **(Figure 1e, f)**. This finding is consistent with previous observations where increased phosphorylation of Orm2 was observed after 2 minutes, followed by its decrease after 10 minutes of heat shock (Sun *et al*, 2012). But we also observed high levels of Orm phosphorylation under basal conditions. In summary, the phosphorylation status of the Orm proteins is difficult to align with the upregulation of SPT after short heat shock. However, the short time frame of increased LCB biosynthesis after heat shock and the importance of serine uptake suggest that a local upregulation of an SPT pool close to the plasma membrane is possible.

### Plasma membrane restricted Ypk1 signaling is sufficient to maintain sphingolipid homeostasis

To further pursue this hypothesis, we tested if peripheral ER restricted TORC2/Ypk1 signaling is sufficient for the Orm dependent heat shock response of yeast cells. Previous studies have indicated that the SPT is differentially regulated in the peripheral ER, where a decrease in Orm2 is observable after myriocin treatment (Breslow *et al*, 2010). The degradation of Orm2 via the EGAD pathway is initiated by phosphorylation from the cytosolic Ypk1 kinase, which is activated in response to signals from the plasma membrane (Schmidt *et al*, 2019; Niles & Powers, 2012; Berchtold *et al*, 2012). To determine if the SPT is mainly regulated in the peripheral ER or if regulation in the entire ER is necessary for Orm regulation, we tethered Ypk1 to the plasma membrane using a CAAX box (Tang *et al*, 2009). Fluorescence microscopy confirmed the tethering **(suppl. Fig. 1a)**. We assessed the functionality of Ypk1-CAAX using genetic interaction studies and tetrad analysis. The deletion of both, *YPK1* and *YPK2,* is lethal, but this lethality could be rescued by the additional deletion of both Orm **(Fig. 2a)** (Roelants *et al*, 2011). *YPK1-CAAX* was also lethal when combined with *ypk2Δ*, but this could be rescued by deleting either *ORM1* or *ORM2* **(suppl. Fig. 1b)**. This indicates that the lethality induced by the deletion of *YPK1/2* is mediated by the dysregulation of SP biosynthesis, although several other targets of Ypk1/2 are known (as previously described by (Roelants *et al*, 2011; Muir *et al*, 2014)).

**Figure 2:**
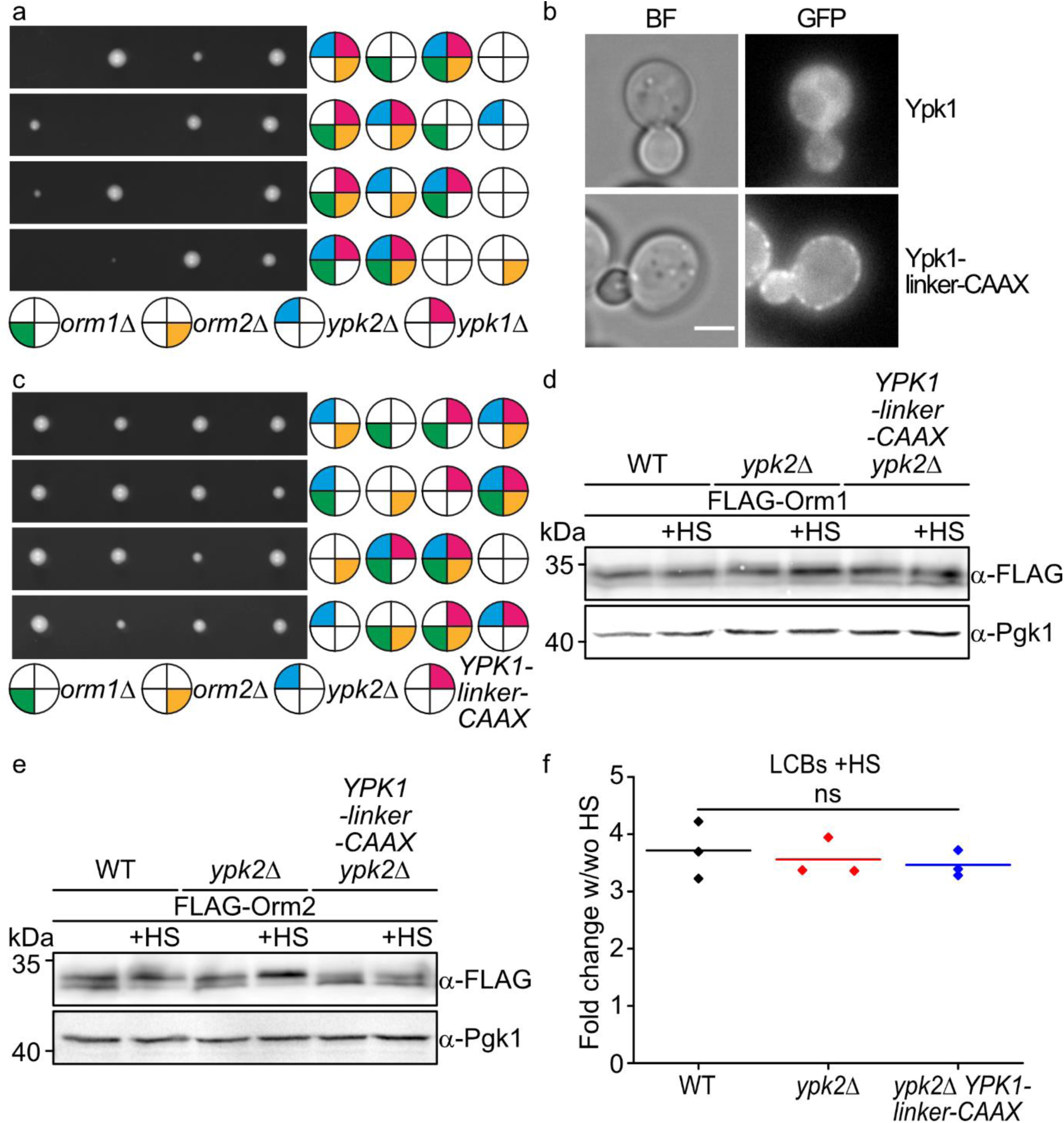
Ypk1 regulates Orm proteins mainly in the peripheral ER. **(a)** Orm proteins are the main target of Ypk1. Tetrad analysis of orm1Δ (green), orm2Δ (yellow), ypk2Δ (blue), and ypk1Δ (pale red) mutants. **(b)** Localization of wildtype GFP-tagged Ypk1 (upper panel) and GFP-linker-CAAX tagged Ypk1 are shown as representative mid-sections. Brightfield images (left panels) and fluorescent images are shown (right panels). Scale bar = 5 µm. **(c)** Plasma membrane targeted Ypk1 with a linker is functional. Tetrad analysis of orm1Δ (green) orm2Δ (yellow) ypk2Δ (blue) and YPK1-linker-CAAX (pale red) cells. **(d,e)** Plasma membrane targeted Ypk1 is only partly able to mediate Orm phosphorylation in response to heat shock. Phosphorylation pattern of **(d)** 3xFLAG-Orm1 and **(e)** 3xFLAG-Orm2 with and without heat shock in WT cells, ypk2Δ cells and ypk2Δ YPK1-linker-CAAX cells. Cells were grown in YPD and subjected to heat shock for 5 minutes at 39°C (+HS) or kept at room temperature. Equal amounts of cells were lysed and analyzed by western blotting using antibodies against the FLAG-tag or Pgk1 as a loading control. **(f)** Plasma membrane targeted Ypk1 is able to upregulate SPT activity. Mass spectrometry-based analysis of LCBs with and without 5 minutes of heat shock at 39° C. Lipids were extracted and analyzed via mass spectrometry. Displayed are the fold changes of LCBs for cells with versus without heat shock treatment in WT, ypk2Δ and ypk2Δ YPK1-linker-CAAX cells. Dots correspond to the values of three independent experiments. *, P-value <0.05, calculated from t-test.

We next added an additional 103 amino acid linker between Ypk1 and the CAAX box to bridge the approximately 30 nm distance between the peripheral ER and the plasma membrane (Gatta *et al*, 2015; Stradalova *et al*, 2012; West *et al*, 2011). This would theoretically allow phosphorylation of the Orm proteins at the peripheral ER from Ypk1-linker-CAAX at the plasma membrane. We confirmed tethering to the plasma membrane with the additional linker using microscopy and tested its functionality using negative genetic interactions with *ypk2Δ* **(Fig. 2b, c)**. *YPK1*-linker-CAAX showed no growth defects when combined with a *YPK2* deletion, demonstrating the functionality of the construct. To exclude major differences induced by the tethering of Ypk1 to the plasma membrane, we also measured the proteome of *YPK1-* linker-CAAX cells. First, we compared *ypk2*Δ cells to WT cells using label-free proteomics. The *YPK2* deletion induced no difference in the proteome, and this was also the case when Ypk1 was tethered to the plasma membrane. The only difference we detected was the decrease of Ypk1 in *ypk2Δ YPK1-linker-CAAX* cells compared to *ypk2Δ* cells or WT cells **(suppl. Fig. 1c, d)**. We also measured phosphorylation of FLAG-tagged Orm1 and Orm2 in the background of *ypk2Δ* and *ypk2Δ YPK1-linker-CAAX* cells. Orm1 phosphorylation, again, remained unchanged under heat shock conditions **(Fig. 2d)**. In contrast, Orm2 phosphorylation was increased in WT cells and *ypk2Δ* cells after heat shock but was largely unaffected in the *ypk2Δ* Ypk1-linker-CAAX strain **(Fig. 2e)**. While these results would indicate that LCB biosynthesis is decreased under these conditions, cells were still able to produce similar amounts of LCBs after heat shock, determined by targeted lipidomics **(Fig. 2f)**. In summary, our results suggest that the currently available tools such as *ORM* deletions, Orm phosphorylation and lipid measurements are insufficient to dissect the spatial and temporal coordination of the rate-limiting step of sphingolipid biosynthesis. We therefore aimed at developing novel tools to analyze SPT regulation and activity with sub-cellular resolution in yeast.

### Developing a nanobody recruitment system to analyze SPT activity with sub-cellular resolution

As mentioned before, we previously suggested that SPT activity is directly coupled to Gnp1 dependent serine uptake at the plasma membrane and Orm2 levels appeared to be especially sensitive to changes in the peripheral ER (Breslow *et al*, 2010; Esch *et al*, 2020). We reasoned that we need tools to isolate the different intracellular pools of the SPT from both the peripheral ER as well as the nuclear ER **(Fig. 3a)**. We developed a nanobody (NB) based recruiting system that depends on a short peptide tag on the SPT as well as a corresponding NB that either targets it to the peripheral ER or to the nuclear ER. The latter is achieved by fusing the NB to either Rtn1 (peripheral ER) or the membrane anchor (amino acids 1-121) of Nvj1 (nuclear ER) (**Fig 3a**) (Millen *et al*, 2008; Kvam & Goldfarb, 2006; Craene *et al*, 2006). As a proof-of-principal experiment, we generated a diploid yeast strain expressing one copy of Lcb1 fused to a GFP and the other copy of Lcb1 fused to mKate and a short ALFA tag (Götzke *et al*, 2019). The two proteins colocalized in both, the peripheral-and the nuclear ER. When we co-expressed a GFP-NB fused to Nvj1_1-121_ and an ALFA-NB fused to Rtn1 we were able to completely separate both pools of Lcb1 **(Fig. 3b)**. While the recruitment was successful we carefully examined if the tagging of the SPT subunits was functional. We therefore, tagged *LCB1* and *LCB2* with either GFP or the short ALFA tag at either the N-terminus or the C-terminus and performed growth tests on control plates or plates containing 0.4 µM myriocin (Wadsworth *et al*, 2013). While all strains grew normally under control conditions, only an N-terminally ALFA tagged *LCB2* strain expressed under the control of its endogenous promotor grew similar to a WT strain on myriocin **(Fig. 3c)**. This suggests that most tags already interfere with the delicate regulation of SPT activity. Based on these results we decided to work in haploid strains recruiting the entire SPT population to either the peripheral or the nuclear ER via the intracellular expressed ALFA-NB (Götzke *et al*, 2019).

**Figure 3:**
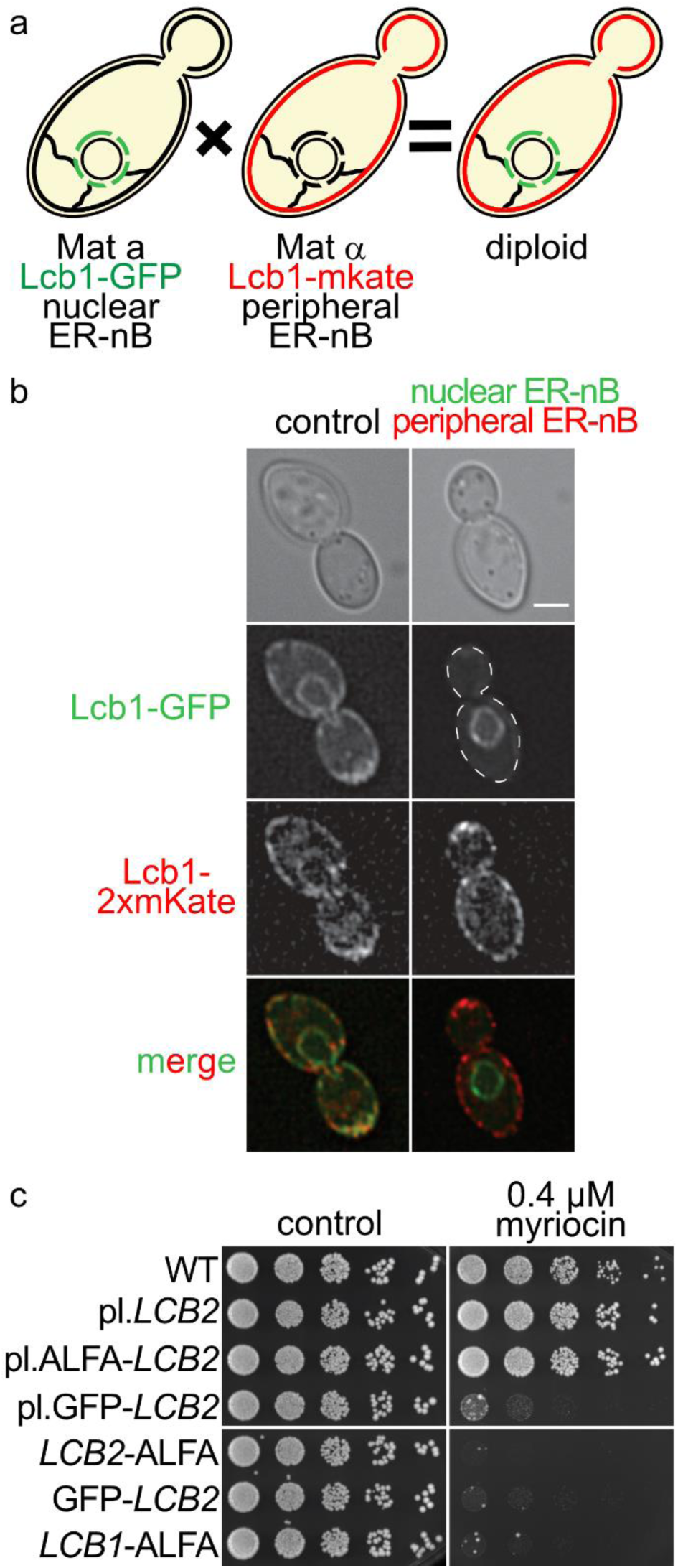
The SPT can be recruited to ER sub-compartments. **(a)** Schematic overview of the diploid strains used in b. **(b)** Localization of GFP-tagged Lcb1 and mkate tagged Lcb1 in a diploid strain either not recruited (left) or recruited to the nuclear and peripheral ER by the use of a nuclear or peripheral ER-nB (right), respectively. Brightfield images (upper panels), GFP signal (upper middle panels) mkate signal (lower middle panels) and merged signals (lower panel) are shown as representative mid-sections. Scale bar = 5 `μ`m. **(c)** Serial dilutions of differentially tagged SPT subunits on YPD plates (control) or plates containing 0.4 `μ`M myriocin. The strains used are from top to bottom: wild-type (WT), lcb2Δ plasmid-LCB2 (SPTallER), plasmid-GFP-LCB2 (pGFP-LCB2), LCB2-ALFA, GFP-LCB2 and LCB1-ALFA

We will refer to cells harboring ALFA-Lcb2 as all ER SPT (SPT^allER^) cells. Cells with ALFA-Lcb2 and the nuclear ER ALFA-NB will be referred to as nuclear ER SPT (SPT^nER^) cells. Cells with ALFA-Lcb2 and a peripheral ER ALFA-NB will be referred to as peripheral ER SPT (SPT^pER^) cells, hereafter **(Fig. 4a)**. First, we utilized the GFP-tagged form of Lcb1, the other catalytically active subunit of the SPT, to confirm recruitment to both parts of the ER. Microscopy confirmed the co-recruitment of the entire population of Lcb1-GFP and thereby also of the ALFA-tagged subunit Lcb2 **(Fig. 4b)**. Next, we analyzed the co-recruitment of the other subunits of the SPOTS complex namely, GFP-Orm1, GFP-Orm2 and GFP-Sac1. Similar to Lcb1, we were also able to co-recruit GFP-Orm1 and GFP-Orm2 to the different ER sub-compartments **(Fig. 4c, d)**. However, we noticed that a small population of GFP-Orm1 and GFP-Orm2 were not co-recruited, suggesting that a pool of Orm proteins exists in the cells that is not bound to the SPOTS complex. In contrast to the other tested proteins, Sac1 recruitment was only possible in the peripheral ER. We did not manage to recruit the Sac1-GFP protein to the nuclear ER via the Nvj1_1-121_-NB fusion protein **(Fig. 4 e)**. This can be either explained by a free pool of Sac1 in the peripheral ER, or by stable protein-protein interactions of Sac1 at the ER-plasma membrane contact site. In line with this observation, Sac1 has been proposed to interact with the VAP proteins (Manford *et al*, 2012). Alternatively, the tag on Lcb2 could interfere with the protein-protein interaction between Lcb2 and Sac1 that is exclusively mediated via its N-terminus (Schäfer *et al*, 2023). To directly test if recruitment interfered with protein-protein interactions we performed co-immunoprecipitations of the recruited SPOTS complex followed by label-free mass spectrometry-based proteomics. Importantly, the affinities of the ALFA-NB with the ALFA-tag are so high that it cannot be used for recruitment and co-immunoprecipitations in parallel. We therefore fused another short peptide tag, the SPOTS tag (Metterlein *et al*, 2018), in front of the ALFA tag. The free SPOTS tag was used for immuno-affinity purification of the SPT^allER^, SPT^nER^ and SPT^pER^. The proteomics analysis revealed that all subunits of the SPOTS complex were co-enriched even in the recruited conditions, suggesting that intra-cellular recruiting does not interfere with the protein-protein interactions within the SPOTS complex. Further, the co-recruitment of the small regulatory subunit Tsc3 was confirmed **(Fig. 5a, b, c)**.

**Figure 4:**
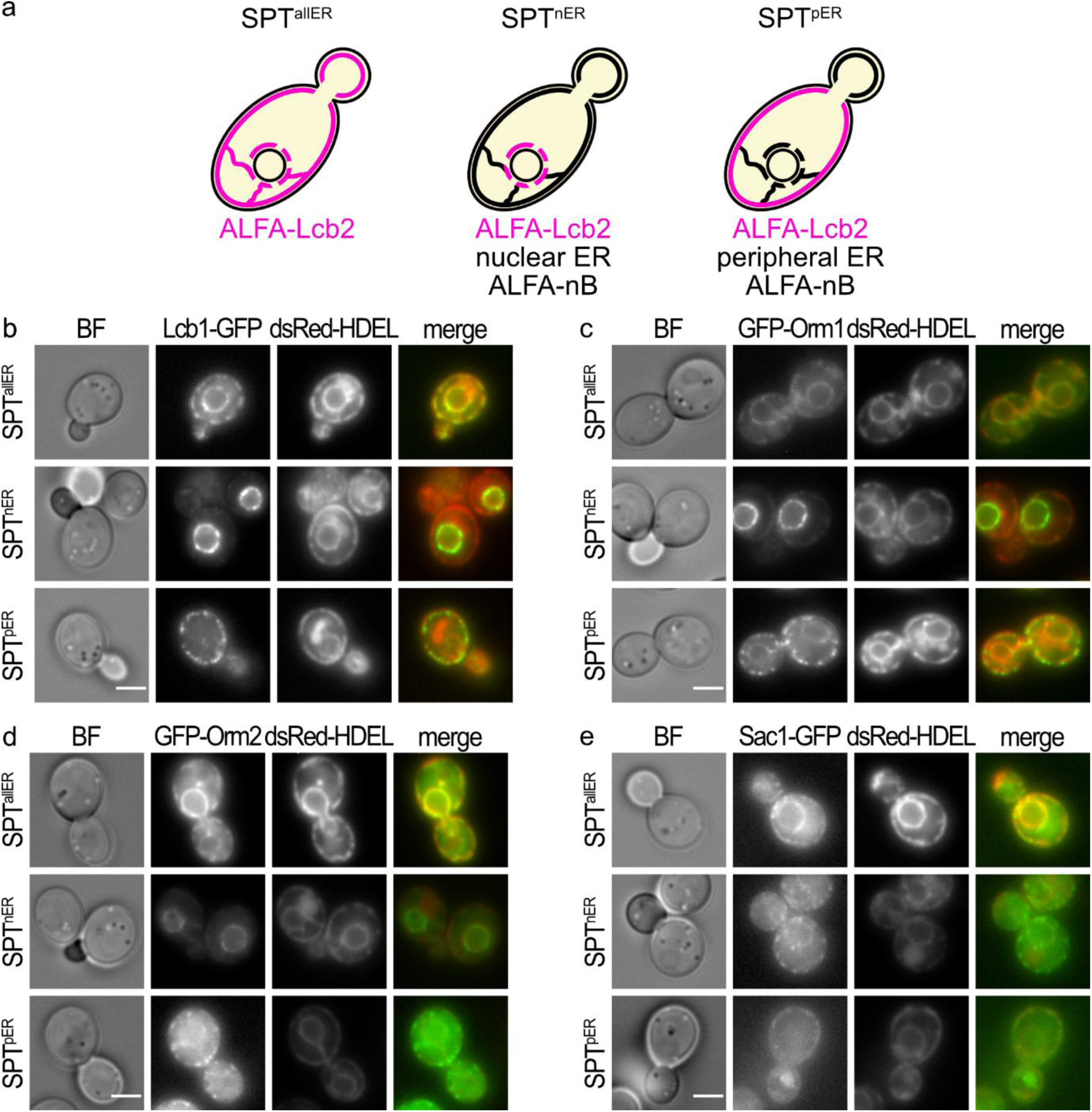
Evaluation of SPOTS complex subunits localization after Lcb2 recruitment of different ER sub-compartments. **(a)** Schematic overview of the system used to rewire the SPT. Cells containing ALFA-tagged Lcb2 will be referred to as all ER SPT (SPT^allER^) cells. Cells containing ALFA-tagged Lcb2 and a nuclear ER ALFA-nB to rewire the SPT to the nuclear ER will be referred to as nuclear ER SPT (SPT^nER^) cells. Cells containing ALFA-tagged Lcb2 and a peripheral ER ALFA-nB to rewire the SPT to the peripheral ER will be referred to as peripheral ER SPT (SPT^pER^) cells. Localization of the SPT within the ER is depicted in pale red. **(b)** Co-localization of Lcb1-GFP in the SPT^allER^ strain (upper panel), the SPT^nER^ strain (middle panel) and in the SPT^pER^ strain (lower panel) with dsRed-HDEL as ER marker are shown as representative mid-sections. Brightfield images (left panels), GFP (middle left panels) and HDEL images (middle right panels) and merged images are shown (right panels). All pictures were processed the same way, except the picture of SPT^nER^ strain for the GFP channel which is shown with lower intensity. Scale bar = 5 µm. **(c)** Co-localization of GFP-Orm1 in the SPT^allER^ strain (upper panel), the SPT^nER^ strain (middle panel) and in the SPT^pER^ strain (lower panel) with dsRed-HDEL as ER marker are shown as representative mid-sections. Brightfield images (left panels), GFP (middle left panels) and HDEL images (middle right panels) and merged images are shown (right panels). All pictures were processed the same way, except the picture of SPT^nER^ strain for the GFP channel which is shown with lower intensity. Scale bar = 5 µm. **(d)** Co-localization of GFP-Orm2 in the SPT^allER^ strain (upper panel), the SPT^nER^ strain (middle panel) and in the SPT^pER^ strain (lower panel) with dsRed-HDEL as ER marker are shown as representative mid-sections. Brightfield images (left panels), GFP (middle left panels) and HDEL images (middle right panels) and merged images are shown (right panels). All pictures were processed the same way, except the picture of SPT^pER^ strain for the GFP channel which is shown with higher intensity. Scale bar = 5 µm. **(e)** Co-localization of Sac1-GFP in the SPT^allER^ strain (upper panel), the SPT^nER^ strain (middle panel) and in the SPT^pER^ strain (lower panel) with dsRed-HDEL as ER marker are shown as representative mid-sections. Brightfield images (left panels), GFP (middle left panels) and HDEL images (middle right panels) and merged images are shown (right panels). All pictures were processed the same way. Scale bar = 5 µm.

**Figure 5:**
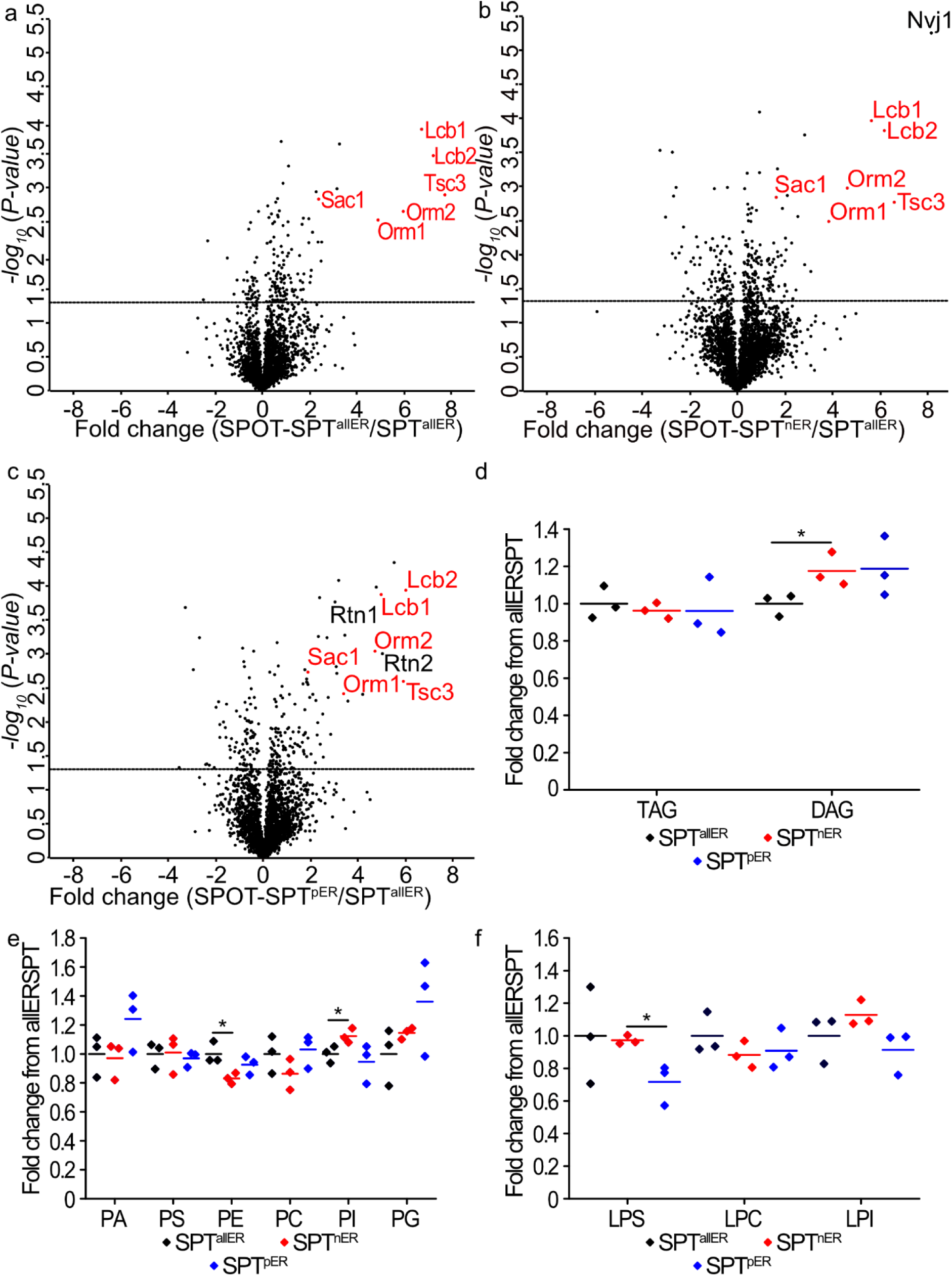
Systematic analysis of the SPT recruited strains by proteomics and lipidomes. The whole SPOTS complex is co-recruited to both parts of the ER. Volcano plots identifying proteins that are co-enriched in SPOT Trap immunoprecipitation experiments. Fold changes were calculated from three independent experiments comparing **(a)** SPOT-SPT^allER^ cells to SPT^allER^ cells, **(b)** SPOT-SPT^nER^ cells to SPT^allER^ cells and **(c)** SPOT-SPT^pER^ cells to SPT^allER^ cells. Fold changes were plotted on the x-axis against negative logarithmic P-values of the t-test performed from three replicates. The dashed line separates significant enriched proteins (P-value <0.05, calculated from t-test) from unaffected proteins. **(d)** The lipidome is mainly unaffected by SPT rewiring. Lipidomic analysis of triacylglycerols (TAG) and diacylglycerols (DAG) of SPT^allER^ cells (black), SPT^nER^ cells (red) and SPT^pER^ cells (blue) are shown as fold changes from SPT^allER^ cells. Dots correspond to the values of three independent experiments. *, P-value <0.05, calculated from t-test. **(e)** Lipidomic analysis of phosphatidic acid (PA), phosphatidylserine (PS), phosphatidylethanolamine (PE), phosphatidylcholine (PC), phosphatidylinositol (PI) and phosphatidylglycerol (PG) of SPT^allER^ cells (black), SPT^nER^ cells (red) and SPT^pER^ cells (blue) are shown as fold changes from SPT^allER^ cells. Dots correspond to the values of three independent experiments. *, P-value <0.05, calculated from t-test. **(f)** Lipidomic analysis of lyso-phosphatidylserine (LPS), lyso-phosphatidylcholine (LPC) and lyso-phosphatidylinositol (LPI) of SPT^allER^ cells (black), SPT^nER^ cells (red) and SPT^pER^ cells (blue) are shown as fold changes from SPT^allER^ cells. Dots correspond to the values of three independent experiments. *, P-value <0.05, calculated from t-test.

To further exclude that the tagging of Lcb2 or SPT recruitment did affect the expression of other proteins, we also analyzed the proteome of the three strains used for recruitment, including SPT^allER^, SPT^nER^, SPT^pER^, and WT cells. No significant changes in protein abundance were observed among the 2945 identified proteins, with minor changes in the expression levels of the SPOTS complex and Rtn1 **(suppl. Fig. 2a-c)**. To control if tagging of Rtn1 affected its function we utilized the negative genetic interactions of *RTN1* with *SPO7* to assess its functionality after peripheral ER rewiring. Spo7 is a component of the Nem1-Spo7 protein phosphatase complex, which controls the function of Pah1 and is essential for triacylglycerol synthesis and nuclear ER morphology (Su *et al*, 2018; Siniossoglou, 1998). Our tetrad analysis showed negative genetic interactions between *spo7Δ* and *rtn1Δ*, whereas *RTN1*-ALFA-nB *spo7Δ* cells displayed normal growth, indicating that tagging Rtn1 for SPT rewiring did not affect its function **(suppl. Fig. 3 d, e).** Additionally, we measured the lipidome in the rewired strains to detect possible differences due to SPT rewiring. We found only minor changes between the three tested strains (SPT^allER^, SPT^nER^, SPT^pER^) **(Fig. 5d, e, f)**. Thus, any potential side effects on SPT activity resulting from changes in the lipidome can be excluded. In summary, the NB based recruitment system works for the SPOTS complex in yeast. As shown here, this system can be utilized to measure protein-protein interactions *in vivo* using fluorescence microscopy as a readout.

### Analysis of ER sub-compartment specific SPOTS complexes

To test if the NB recruitment system allows to detect changes in the activity of the different SPT pools, we measured LCB levels in the SPT^allER^, SPT^nER^ and SPT^pER^ strains using targeted lipidomics under control and heat shock conditions. LCBs increase after heat shock was not significantly changed in response to cellular SPT localization **(Fig. 6a)**. We also analyzed phosphorylation of FLAG-Orm1 and FLAG-Orm2 in the strains with differentially recruited SPTs. This analysis showed increased phosphorylation levels of Orm2 at all localizations after heat shock, again with relative high levels of basal phosphorylation represented by the slower running band **(Fig. 6b, c)**. Nevertheless, we cannot differentiate whether the Ypk1 kinase can phosphorylate the Orm proteins within the SPOTS complexes in both cellular localizations or if the free Orm protein pool is sufficient to achieve the observed elevated levels of Orm phosphorylation. Next, we investigated if the deletion of either of the *ORM* genes had different effects in the SPT recruited strains in control and heat shock conditions. Targeted lipidomics revealed that the deletion of *ORM1* had only a minimal effect on LCB levels in the SPT^allER^ strain and the SPT^pER^ strain but let to a small but significant increase in LCB levels in the SPT^nER^ strain **(Fig. 6d)**. In contrast, deletion of *ORM2* already led to a nearly 4-fold increase in LCB levels in the SPT^allER^ strain and the SPT^pER^ strains and to a smaller increase in the SPT^nER^ strain **(Fig. 6e)**. Exposing the cells to 5 minutes of heat shock led to a 5-fold increase in LCB levels in all *orm1Δ* strains **(Fig. 6e)**. In contrast, exposing *orm2Δ* strains to heat shock conditions only resulted in a small increase in LCB levels under all conditions **(Fig. 6f)**. Together, we find small differences in the activities of the recruited SPT strains depending on the overall localization of the SPT together with their inhibition by Orm1 and Orm2.

**Figure 6:**
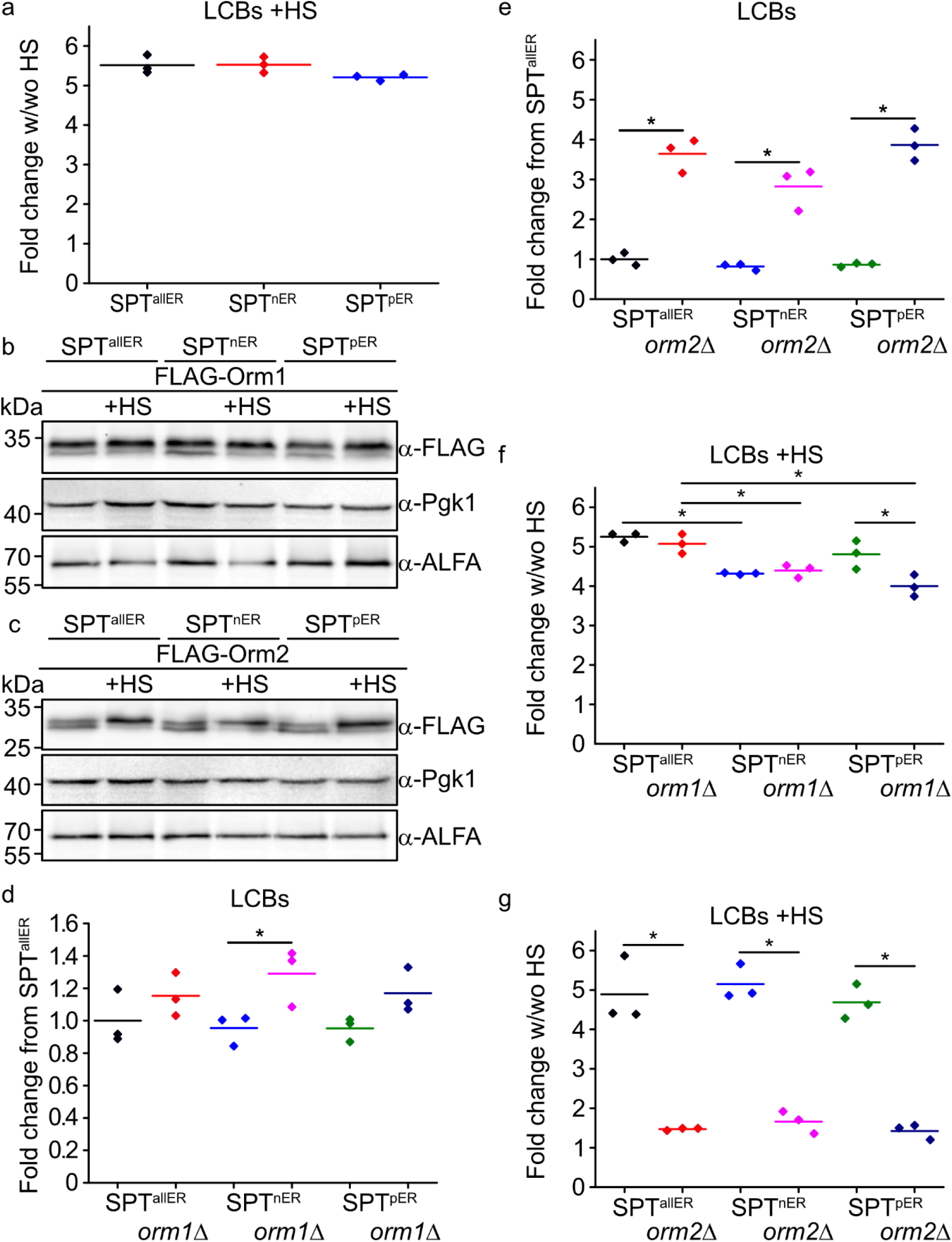
Determining SPOTS activity in different sub-compartments. **(a).** Mass spectrometry-based analysis of LCBs with and without 5 minutes of heat shock at 39° C. Displayed are the fold changes of LCBs for cells with versus without heat shock treatment in SPT^allER^ (black), SPT^nER^ (red) and SPT^pER^ (blue) strains. Dots correspond to the values of three independent experiments. *, P-value <0.05, calculated from t-test. **(b,c)** Phosphorylation pattern of **(b)** 3xFLAG-Orm1 and **(c)** 3xFLAG-Orm2 with and without heat shock in SPT^allER^ cells, SPT^nER^ cells and SPT^pER^ cells. Cells were grown in YPD and subjected to heat shock for 5 minutes at 39°C (+HS) or kept at room temperature. Equal amounts of cells were lysed and analyzed by western blotting using antibodies against the FLAG-tag or Pgk1 as a loading control. **(d)** Orm1 inhibits the SPT more in the nuclear ER. Cells were grown in YPD medium. Lipids were extracted and analyzed via mass spectrometry. Displayed are the amounts of total LCBs as fold change from SPT^allER^ cells. The strains used are: SPT^allER^ (black) cells, SPT^allER^ orm1Δ (red) cells, SPT^nER^ (blue) cells, SPT^nER^ orm1Δ (pale red) cells, SPT^pER^ (green) cells and SPT^pER^ orm1Δ (dark blue) cells. Dots correspond to the values of three independent experiments. *, P-value <0.05, calculated from t-test. **(e)** Orm2 inhibits the SPT less strong in the nuclear ER. Cells were grown in YPD medium. Lipids were extracted and analyzed via mass spectrometry. Displayed are the amounts of total LCBs as fold change from SPT^allER^ cells. The strains used are: SPT^allER^ (black) cells, SPT^allER^ orm2Δ (red) cells, SPT^nER^ (blue) cells, SPT^nER^ orm2Δ (pale red) cells, SPT^pER^ (green) cells and SPT^pER^ orm2Δ (dark blue) cells each without (upper plot) and with heat shock (lower plot). Dots correspond to the values of three independent experiments. *, P-value <0.05, calculated from t-test. **(f)** Mass spectrometry-based analysis of LCBs with and without 5 minutes of heat shock at 39° C. Displayed are the fold changes of LCBs for cells with versus without heat shock treatment. The strains used are: SPT^allER^ (black) cells, SPT^allER^ orm1Δ (red) cells, SPT^nER^ (blue) cells, SPT^nER^ orm1Δ (pale red) cells, SPT^pER^ (green) cells and SPT^pER^ orm1Δ (dark blue) cells. Dots correspond to the values of three independent experiments. *, P-value <0.05, calculated from t-test. **(g)** Mass spectrometry-based analysis of LCBs with and without 5 minutes of heat shock at 39° C. Displayed are the fold changes of LCBs for cells with versus without heat shock treatment. The strains used are: SPT^allER^ (black) cells, SPT^allER^ orm2Δ (red) cells, SPT^nER^ (blue) cells, SPT^nER^ orm2Δ (pale red) cells, SPT^pER^ (green) cells and SPT^pER^ orm2Δ (dark blue) cells. Dots correspond to the values of three independent experiments. *, P-value <0.05, calculated from t-test.

### Combining SPOTS recruitment with serine pulse labeling allows the detection of differentially active SPT pools

To test if the lack of differences in the LCB levels in the analyzed strains is based on the lack of sensitivity of our detection method we used pulse labeling of LCBs with ^13^C_31_^5^N_1_-serine (Martínez-Montañés *et al*, 2020; Esch *et al*, 2020). During the SPT catalyzed condensation of serine and palmitoyl-CoA carbon dioxide from the serine is lost. Therefore, we expect a mass difference of +3 for all ^13^C_3_^15^N_1_-serine labeled sphingolipids **(Fig. 7a)**. We added labeled serine and followed its incorporation into LCBs and ceramide over a time course of 30 mins in SPT^allER^, SPT^nER^ and SPT^pER^ strains. This analysis showed that the incorporation of serine was slower in the SPT^nER^ strains compared to the SPT^allER^ and SPT^pER^ strains, suggesting that the activity of the SPT is lower when recruited to the nuclear ER **(Fig. 7b)**. The same effect was observed with a 10 min delay in the levels of labelled ceramides **(Fig. 7c)**. Interestingly, the overall levels of LCBs and ceramides remained largely unchanged under the used conditions **(Fig. 7b, c)**. When we exposed the cells to heat shock and measured the incorporation of labeled serine into LCBs and ceramides, we observed comparable incorporation rates in the SPT^nER^ strain **(Fig. 7d, e)**. However, incorporation of labelled LCBs into ceramides was reduced in the SPT^nER^ cells after heat shock **(Fig. 7f)**. Thus, decreased SPT^nER^ activity seems to result in an overall decreased flux through the sphingolipid biosynthesis pathway. In summary, combining the NB recruitment system with pulse labeling experiments allows the detection of altered activities of the SPT pools in different sub-cellular compartments.

**Figure 7:**
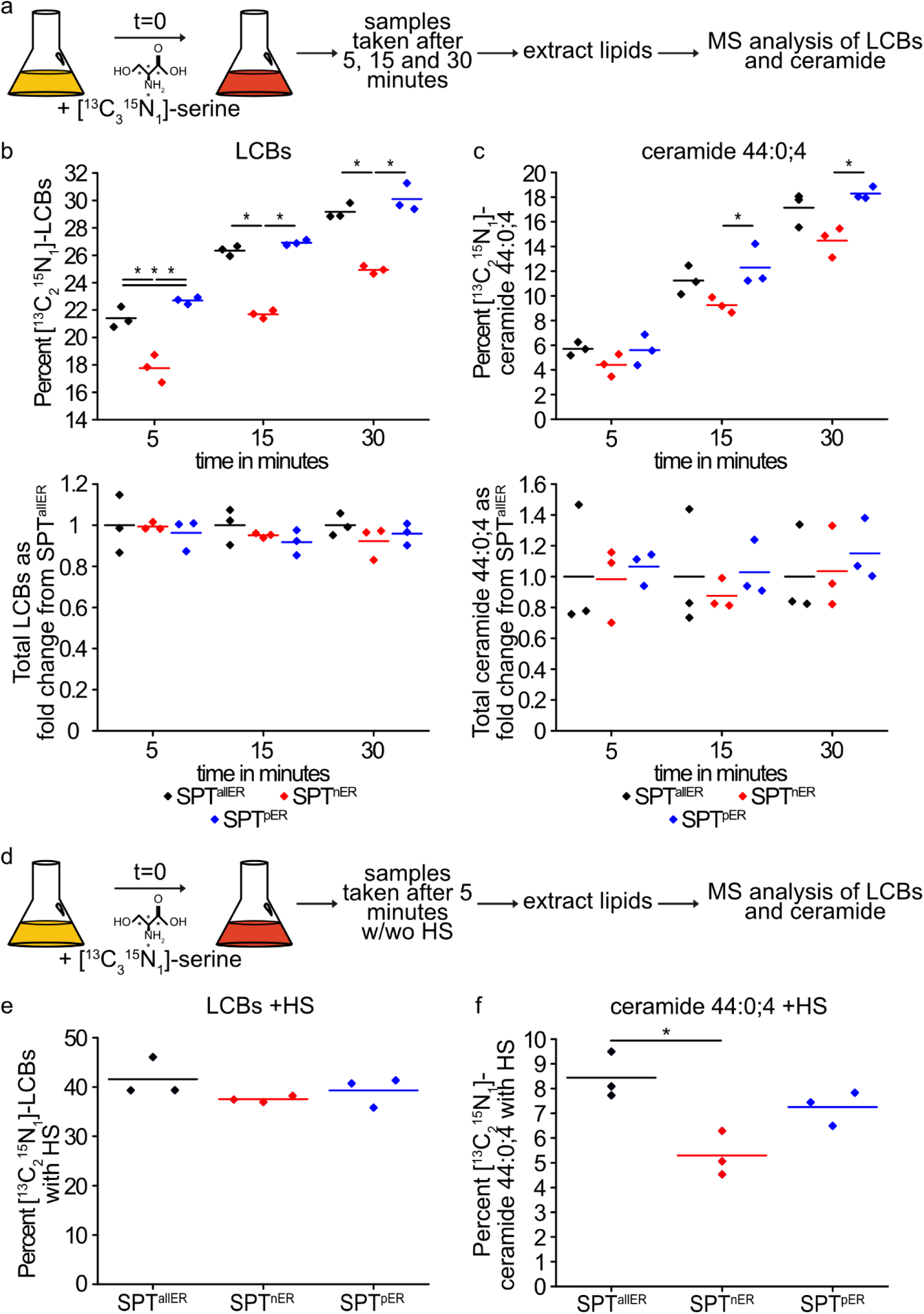
SPT activity is changed in ER sub-compartments. **(a)** Schematic overview of the experimental setup for the performed flux analysis shown in b and c. **(b)** Integration of [^13^C3^15^N1]-serine into long chain bases (LCBs). Cells were labelled with [^13^C3^15^N1]-serine for 5, 15 and 30 minutes in YPD medium. Displayed are the amounts of [^13^C3^15^N1]-serine labelled LCBs as percent of total LCBs (upper panel) and total LCBs (lower panel) of as fold change from SPT^allER^ cells. The strains used are: SPT^allER^ (black), SPT^nER^ (red) and SPT^pER^ (blue) cells. Dots correspond to the values of three independent experiments. *, P-value <0.05, calculated from t-test. **(c)** Integration of [^13^C3^15^N1]-serine into ceramides. Cells were labelled with [^13^C3^15^N1]-serine for 5, 15 and 30 minutes in YPD medium. Lipids were extracted and analyzed via mass spectrometry. Displayed are the amounts of [^13^C3^15^N1]-serine labelled 44:0;4 ceramide as percent of total 44:0;4 ceramide (upper panel) and total 44:0;4 ceramide (lower panel) as fold change from SPT^allER^ cells. The strains used are: of SPT^allER^ (black), SPT^nER^ (red) and SPT^pER^ (blue) cells. Dots correspond to the values of three independent experiments.*, P-value <0.05, calculated from t-test. **(d)** Schematic overview of the experimental setup for the performed [^13^C3^15^N1]-serine labelling with 5 minutes of heat shock shown in d and e. **(e)** Integration of [^13^C3^15^N1]-serine into long chain bases (LCBs) after heat shock. Cells were labelled with [^13^C3^15^N1]-serine for 5 minutes in YPD medium with or without heat shock. Displayed are the amounts of heat shock treated [^13^C3^15^N1]-serine labelled LCBs as percent of total LCBs. The strains used are: SPT^allER^ (black), SPT^nER^ (red) and SPT^pER^ (blue) cells. Dots correspond to the values of three independent experiments. *, P-value <0.05, calculated from t-test. **(f)** Integration of [^13^C3^15^N1]-serine into ceramides after heat shock. Cells were labelled with [^13^C3^15^N1]-serine for 5 minutes in YPD medium with or without heat shock. Lipids were extracted and analyzed via mass spectrometry. Displayed are the amounts of heat shock treated [^13^C3^15^N1]-serine labelled ceramides 44:0;4 as percent of total ceramides 44:0;4. The strains used are: SPT^allER^ (black), SPT^nER^ (red) and SPT^pER^ (blue) cells. Dots correspond to the values of three independent experiments. *, P-value <0.05, calculated from t-test.

## Discussion

The condensation of serine and palmitoyl-CoA catalyzed by the SPT is the rate limiting step in sphingolipid biosynthesis. It is clear that multiple input signals control this step, for example phosphorylation of the Orm proteins by the TORC2 signaling pathway as well as the levels of the downstream metabolite ceramide (Davis *et al*, 2019; Roelants *et al*, 2011; Breslow *et al*, 2010; Berchtold *et al*, 2012; Niles & Powers, 2012). Besides this already complex regulatory network, we previously suggested differentially regulated pools of the SPT in the nuclear and the peripheral ER (Esch *et al*, 2020). However, determining the activities of the same enzyme in different sub-cellular compartments is extremely challenging. Here we used SPT as a model to develop a system that is able to determine lipid metabolic enzyme activities in different parts of the ER. We combined a NB based recruitment system with a pulse labelling approach to analyze the activity and regulation of different SPT pools in yeast cells. Together, the system we developed here is a first step towards studying lipid metabolism with sub-cellular resolution. Intracellular expressed NBs combined with small complementing peptide tags have previously been used to alter the localization of proteins in the cell (Traenkle *et al*, 2020). Similarly, the rapamycin induced targeting system represents an inducible re-localization approach (Chen *et al*, 1995). Our analysis of the SPT recruitment system to different ER sub-compartments shows that the system can be used to recruit entire protein complexes. Recruiting one subunit of a complex allows the co-recruitment of the other subunits and therefore can also be used as a system to measure protein-protein interactions *in vivo*, using fluorescence microscopy as a readout. Similar assays have been established for example in mammalian cells (Sotolongo Bellón *et al*, 2022). Interestingly, it was only possible to completely recruit the Sac1 subunit of the SPOTS complex to the peripheral ER and not to the nuclear ER, suggesting that a non-SPT bound Sac1 pool only localizes to the peripheral ER. In line with this observation, Sac1 was previously shown to interact with other proteins in the cell including the VAP proteins that form pER-plasma membrane contacts (Manford *et al*, 2012).

Combining the SPT recruitment system with pulse labeling of sphingolipid intermediates allows the detection of different SPT pools in the cell. So far it was only possible to measure bulk activity of lipid metabolic enzymes in the cell. Other complementary approaches are, for example, lipidomics MALDI imaging mass spectrometry (MALDI-IMS) that allows the detection of lipids in a certain cellular sub-compartment (Dreisewerd *et al*, 2022; Soltwisch *et al*, 2015). The resolution of this approach is limited and therefore will be difficult to adapt to yeast cells. Other assays that allow the detection of local lipid biosynthesis, modification and lipid transport were recently shown by the Kornmann lab (John Peter *et al*, 2022a, 2022b). Here, yeast cells express non-yeast lipid modifying enzymes in different sub-compartments that allow the visualization of lipid biosynthesis and transport.

Why is it important to measure lipid metabolism with sub-cellular resolution? A prominent example are the activities of the phosphatidylserine decarboxylases (Psd) Psd1 and Psd2 in yeast cells. Both enzymes catalyze the same reaction, the decarboxylation of phosphatidylserine to phosphatidylethanolamine (Voelker, 1997). While Psd1 is described as a mitochondrial and ER localized enzyme, Psd2 is described as both endosomal and Golgi localized (Friedman *et al*, 2018; Gulshan *et al*, 2010). Approaches to target Psd1 to either mitochondria or the ER based on different targeting sequences have already been developed (Friedman *et al*, 2018). Our approach would allow similar experiments for proteins that cannot be targeted just by organelle targeting motifs. Another prominent example for the compartmentation of a membrane is the yeast the plasma membrane. Here, amino acid transporter activity depends on their localization in and out of a specialized domain formed by eisosomes (Gournas *et al*, 2018; Busto *et al*, 2018; Walther *et al*, 2006). Other examples are the differences between the tubular ER and ER sheets in mammalian cells. Are certain lipid metabolic enzymes only active in one of the compartments? Other possible applications could be the analysis of entire metabolic pathways and enzyme super complexes formation analogous to the mitochondrial super-complexes (Robinson & Srere, 1985). Using yeast sphingolipid metabolism as an example one could imagine to recruit the SPT to the nuclear ER while the subsequent enzyme Tsc10 is recruited to the peripheral ER. This would allow the determination of the need for substrate handover. These questions could be addressed using a similar approach as presented here.

The used approach also has its limitations that should be addressed in the future. The recruitment of the entire SPT population to one sub-compartment changes the overall enzyme amount at this location. This by itself could already lead to changes in the activity. It also remains possible that oligomerization of the SPT complexes regulates their activity in the different ER sub-compartments (Hornemann *et al*, 2007; Li *et al*, 2021; Wang *et al*, 2021; Han *et al*, 2019), which may be susceptible to local changes in enzyme abundance. In addition, the non-inducible recruitment could allow the cells to adapt to the changing conditions and therefore modulate SPT activity by homeostatic regulations. Thus, an inducible recruitment system would be even more preferable.

In case of the complex SPT regulation network the recruitment system helped us to gain novel insights. We indeed find two differentially active SPT pools in the cells with the less active one at the nuclear ER. This would be in line with our previous hypothesis that serine taken up by the cells is preferentially incorporated into LCBs at the peripheral ER (Esch *et al*, 2020). It also appears that the Orm2 protein is more important to control the heat shock induced increase in LCBs in yeast cells based on our results. In a recent study we have solved the structure of the Orm1 containing SPOTS complex and have proposed that a monomeric form is the predominant form in yeast cells (Schäfer *et al*, 2023). This could also explain why the two Orm proteins are differentially regulated by the EGAD pathway (Schmidt *et al*, 2019; Bhaduri *et al*, 2023). It is also noteworthy that the timing of the increase in LCB biosynthesis after 5 minutes of heat shock and the regulatory mechanism by the EGAD pathway are difficult to align. Therefore, we suspect that the phosphorylation of the Orm proteins must either lead to a conformational change allowing the release of bound ceramide or directly lead to the dissociation of the Orm protein from the SPOTS complex (Schäfer *et al*, 2023; Davis *et al*, 2019). Taken together, our results shed some new light on the already complex regulatory network of the SPT but will require further investigations. The here presented combination of NB recruitment of lipid metabolic enzymes combined with pulse labeling approaches is one further step to tackle this complex regulatory system and can also be used to study other important lipid metabolic pathways.

## Materials and Methods

### Yeast strains, plasmids, and media

Yeast strains used in this study are shown in Table1. All deletions and tagging of all proteins were performed as described in (Janke *et al*, 2004). All plasmids used in this study are shown in Table2. All oligonucleotides used in this study are shown in Table3. Sequences were cloned into plasmid vectors via fast cloning (Li *et al*, 2011) and the ALFA tag, SPOT tag, CAAX box and 103aa linker were inserted using Q5 mutagenesis (Götzke *et al*, 2019; Tang *et al*, 2009; Gatta *et al*, 2015; Metterlein *et al*, 2018). All experiments were performed in normal YPD media. For microscopy experiments SDC-lysine media was used (2% glucose, 6.75 g/l yeast nitrogen base without amino acids (291929, BD Difco), 1.92 g/l yeast synthetic drop-out media supplements without lysine (Y1896; Sigma Aldrich) supplemented with 30 mg/ml lysine. Sporulation plates were made with 1 % potassium acetate and 3 % agar.

**Table 1:**
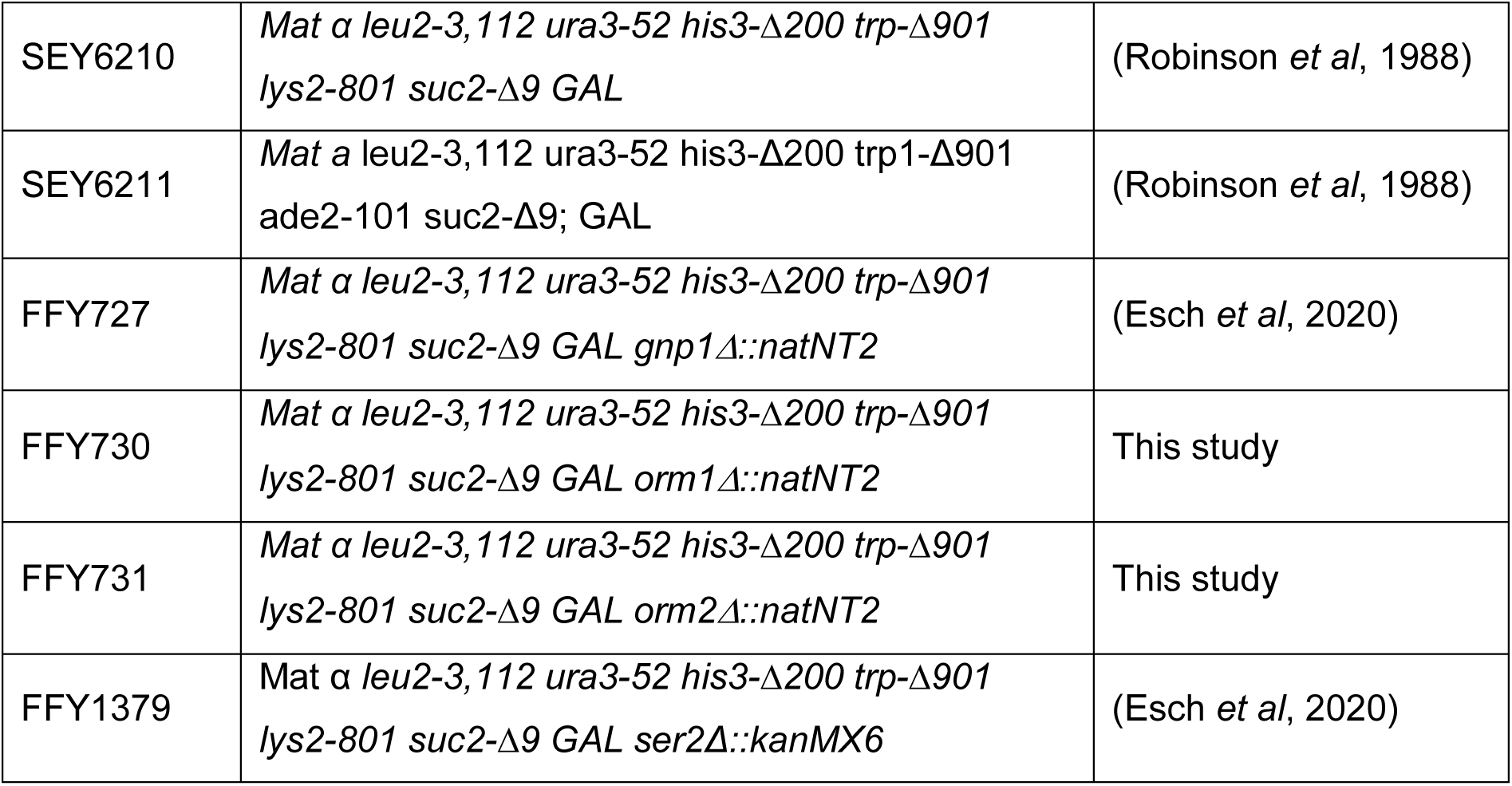
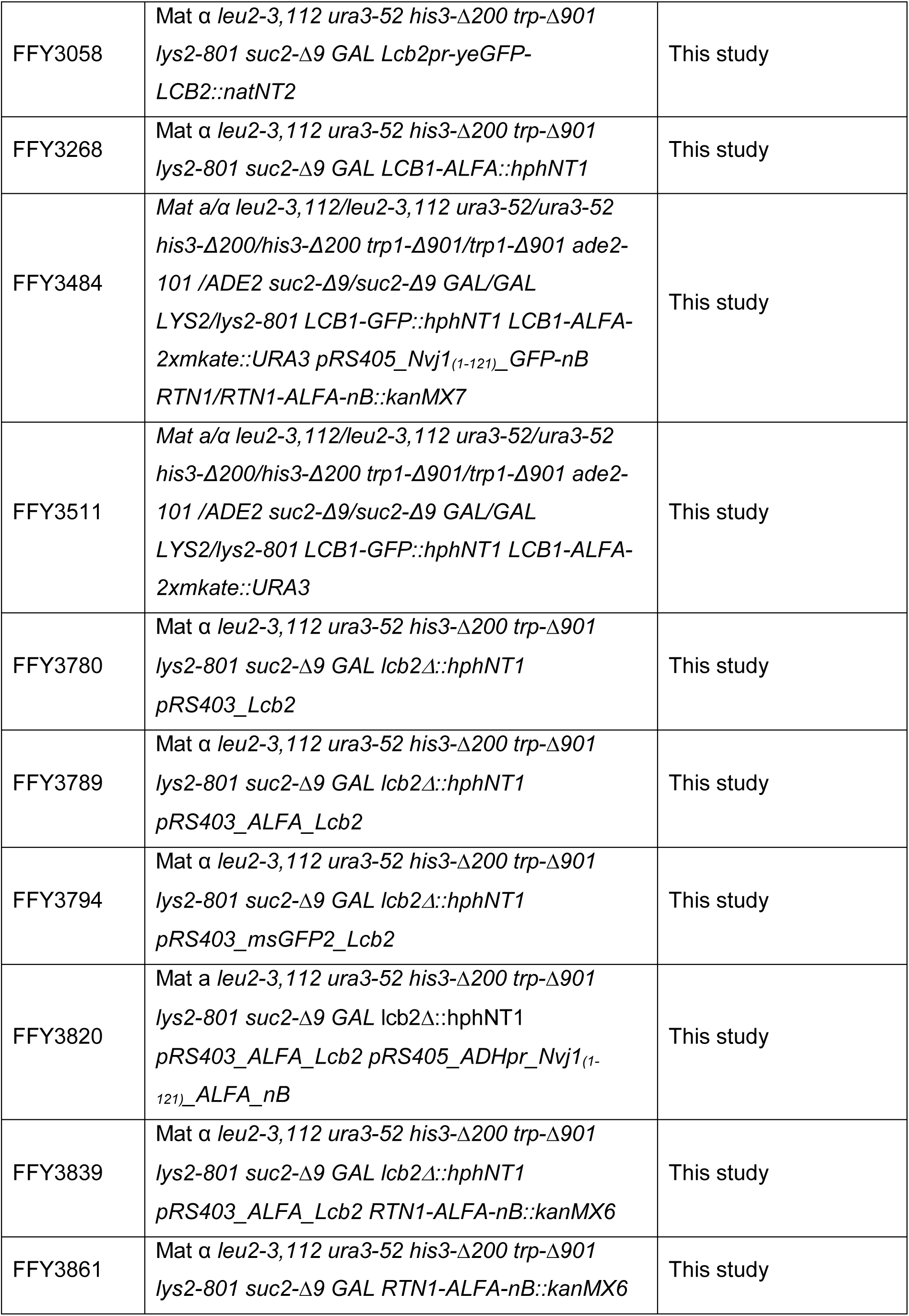
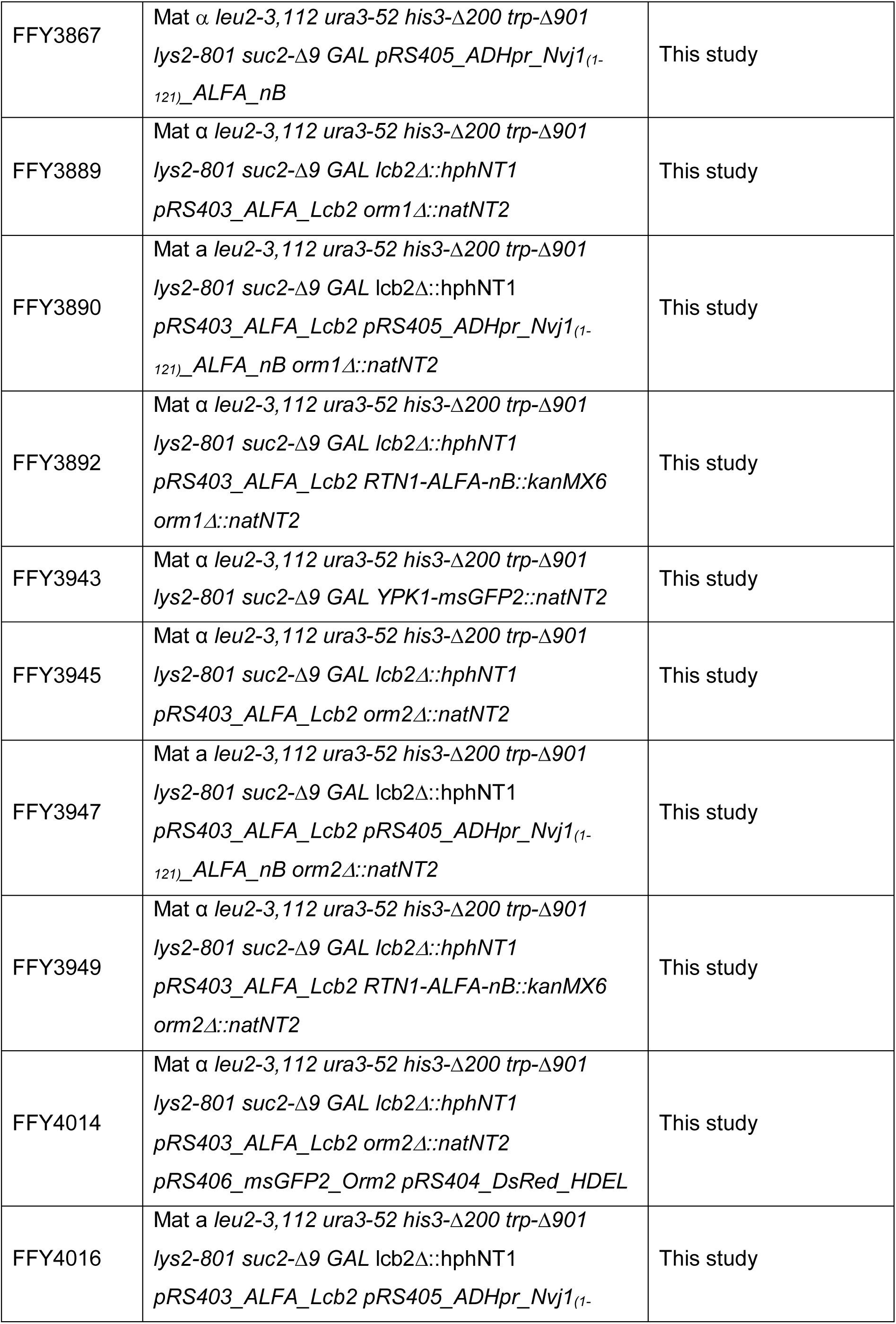
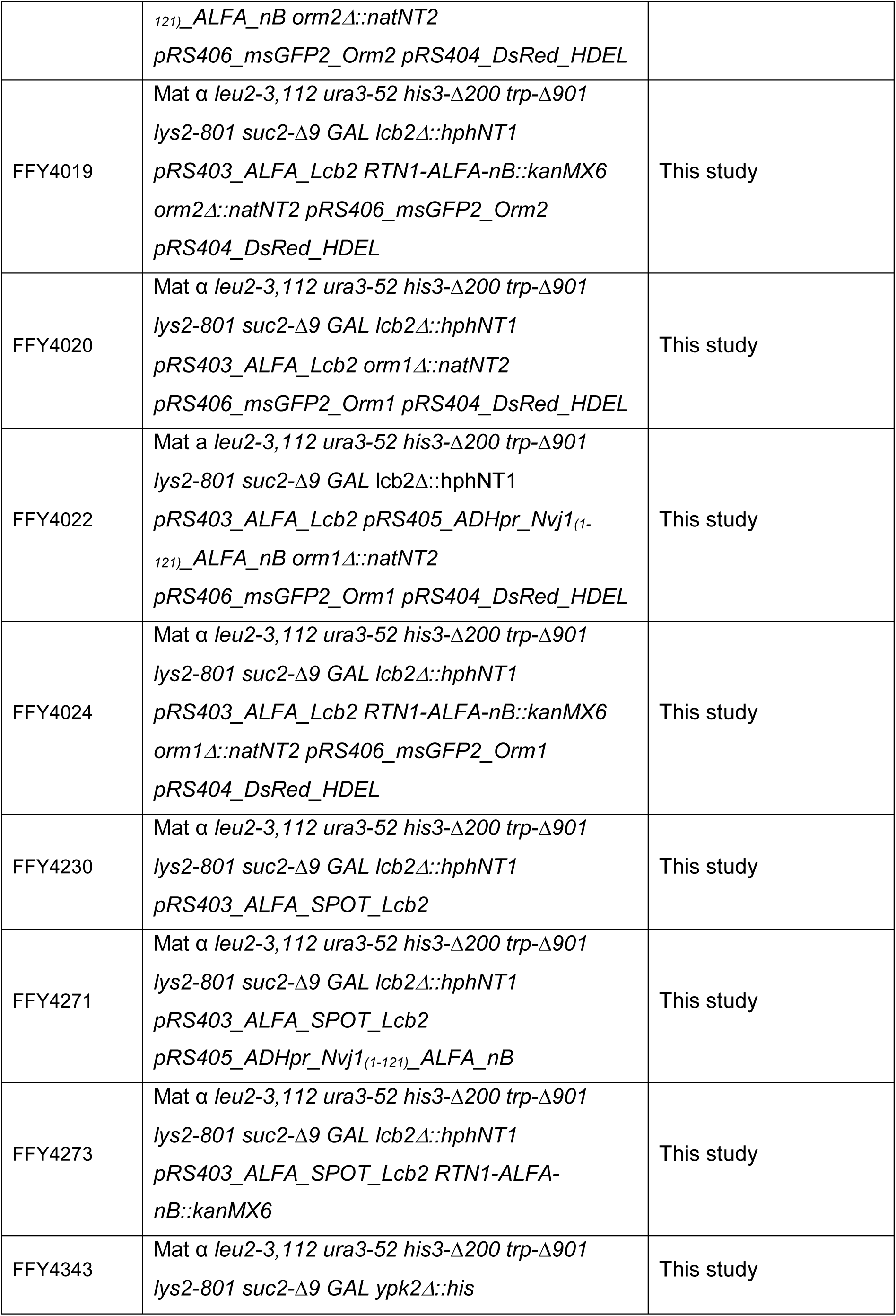
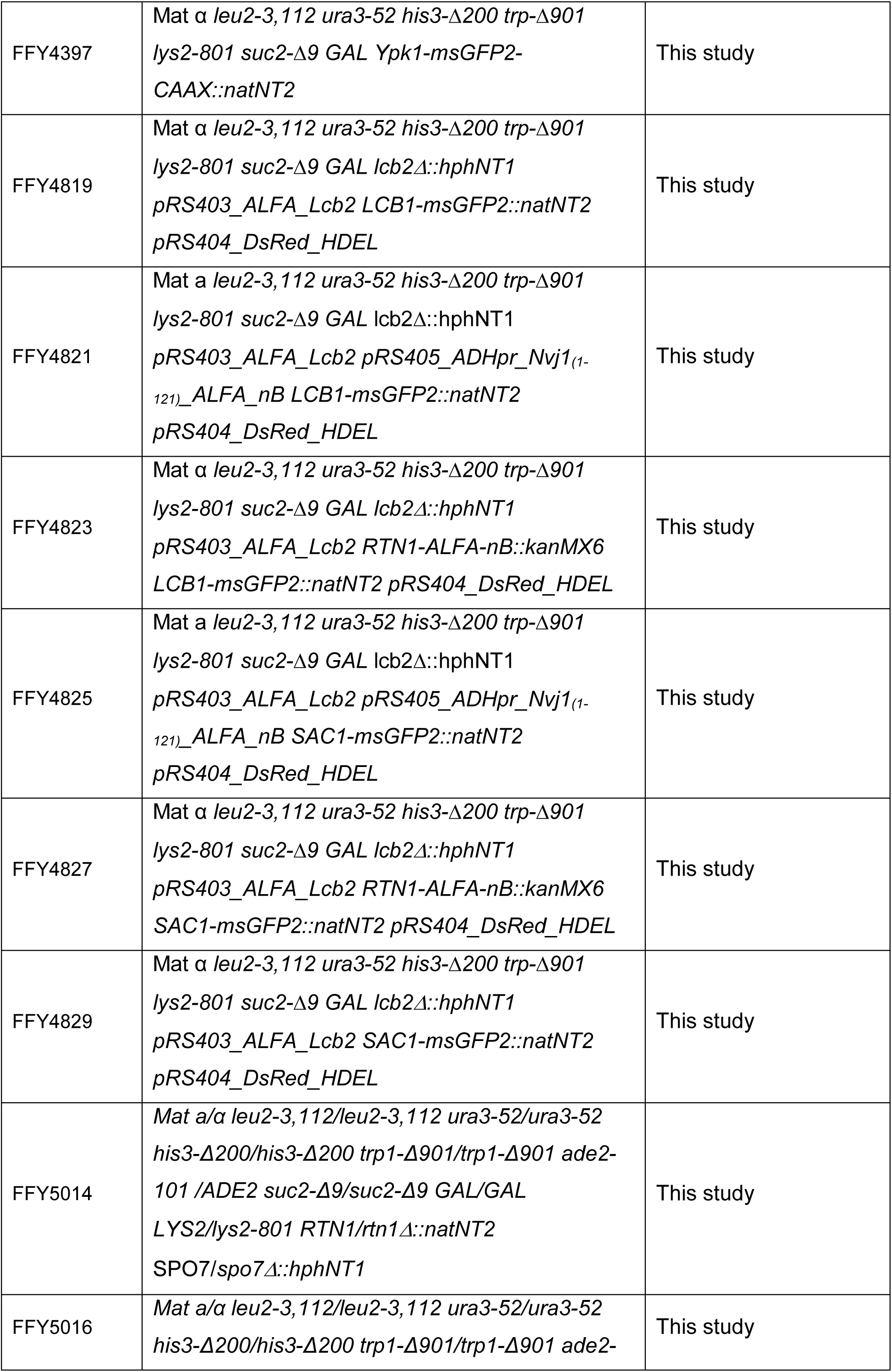
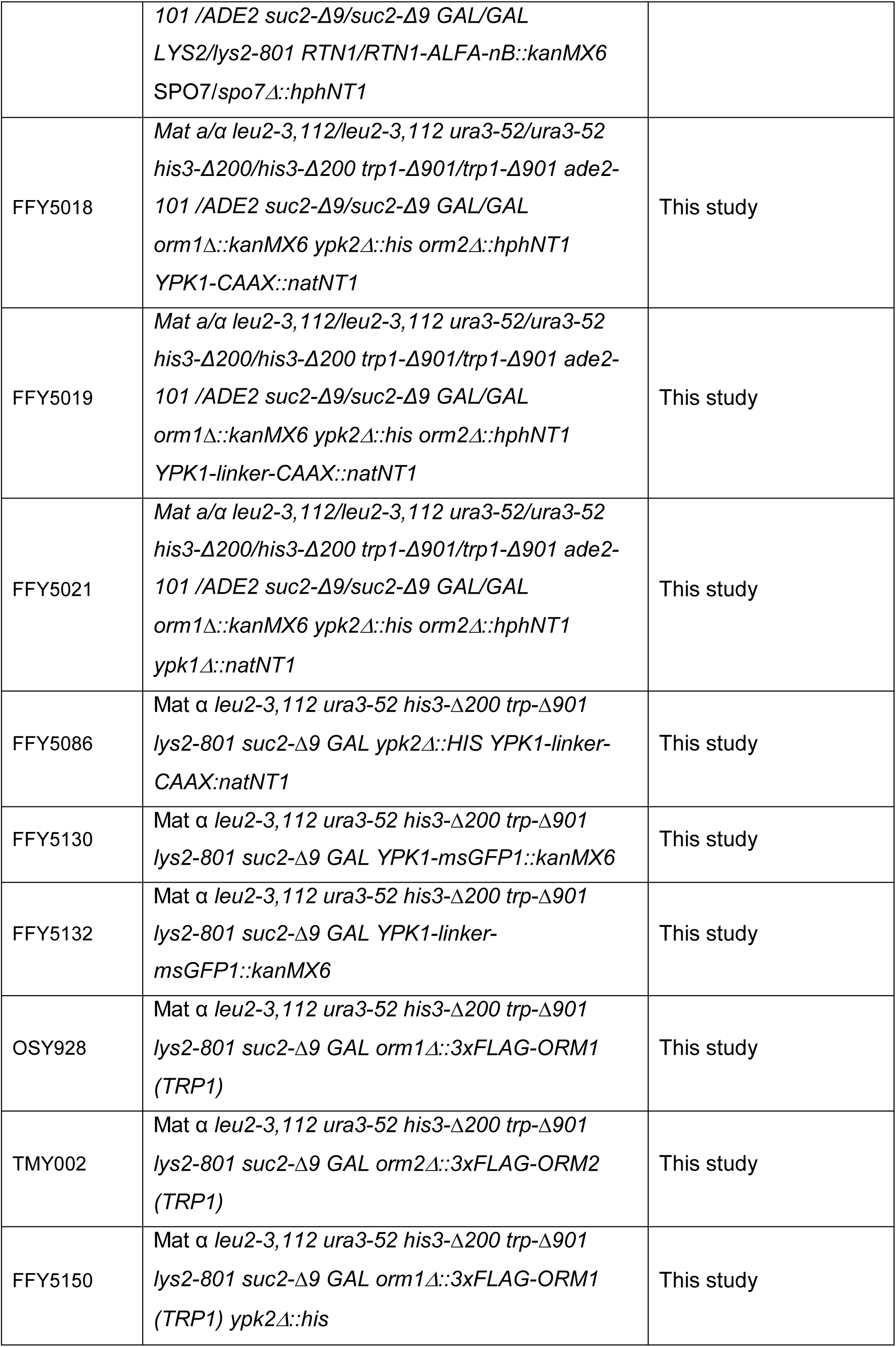
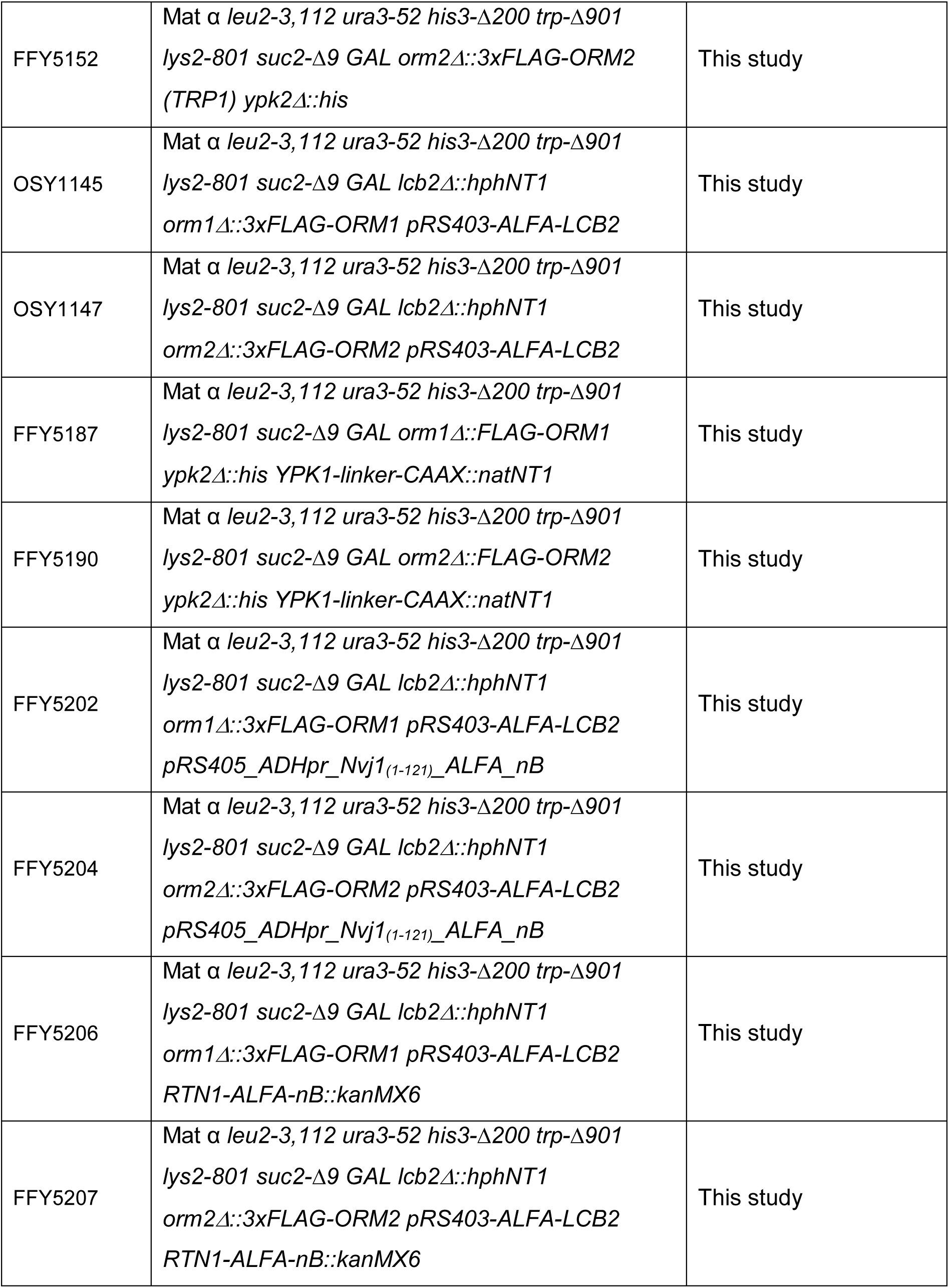
List of all yeast strains and their genotypes used in this study.

**Table 2:** List of all plasmids used in this study.

|  |  |  |
| --- | --- | --- |
| FFP313 | pRS405-ADHpr-NVJ1<sub>(1-121)</sub>-GFP-nB | This study |
| FFP330 | pYM-N9-LCB2prom-yeGFP | This study |
| FFP349 | pYM40-ALFA-tag | This study |
| FFP352 | pYM38-ALFA-nB | This study |
| FFP368 | pRS403-Lcb2 | This study |
| FFP386 | pRS405-ADHpr-NVJ1<sub>(1-121)</sub>-ALFA-nB | This study |
| FFP410 | pRS403-ALFA-Lcb2 | (Schäfer *et al*, 2023) |
| FFP416 | pRS403-msGFP2-LCB2 | This study |
| FFP425 | pRS404-DsRed-HDEL | (Schäfer *et al*, 2023) |
| FFP451 | pRS406-msGFP2-Orm2 | This study |
| FFP459 | pRS406-GFP-Orm1 | This study |
| FFP492 | pRS403-ALFA-SPOT-Lcb2 | This study |
| FFP525 | pYM-msGFP2-CAAX | This study |
| FFP528 | pYM-CAAX | This study |
| FFP581 | pYM-linker-CAAX | This study |
| pOS265 | pFA6a-Orm1pr-3xFLAG-Orm1-3'Orm1 | This study |
| pTM001 | pFA6a-Orm2pr-3xFLAG-Orm2-3'Orm2 | This study |
|  | pYM-msGFP1 | Ungermann Lab |
|  | pYM-msGFP2 | Ungermann Lab |
|  | pYM-yeGFP | Ungermann Lab |
|  | pYM-2xmkate | Ungermann Lab |

**Table 3:**
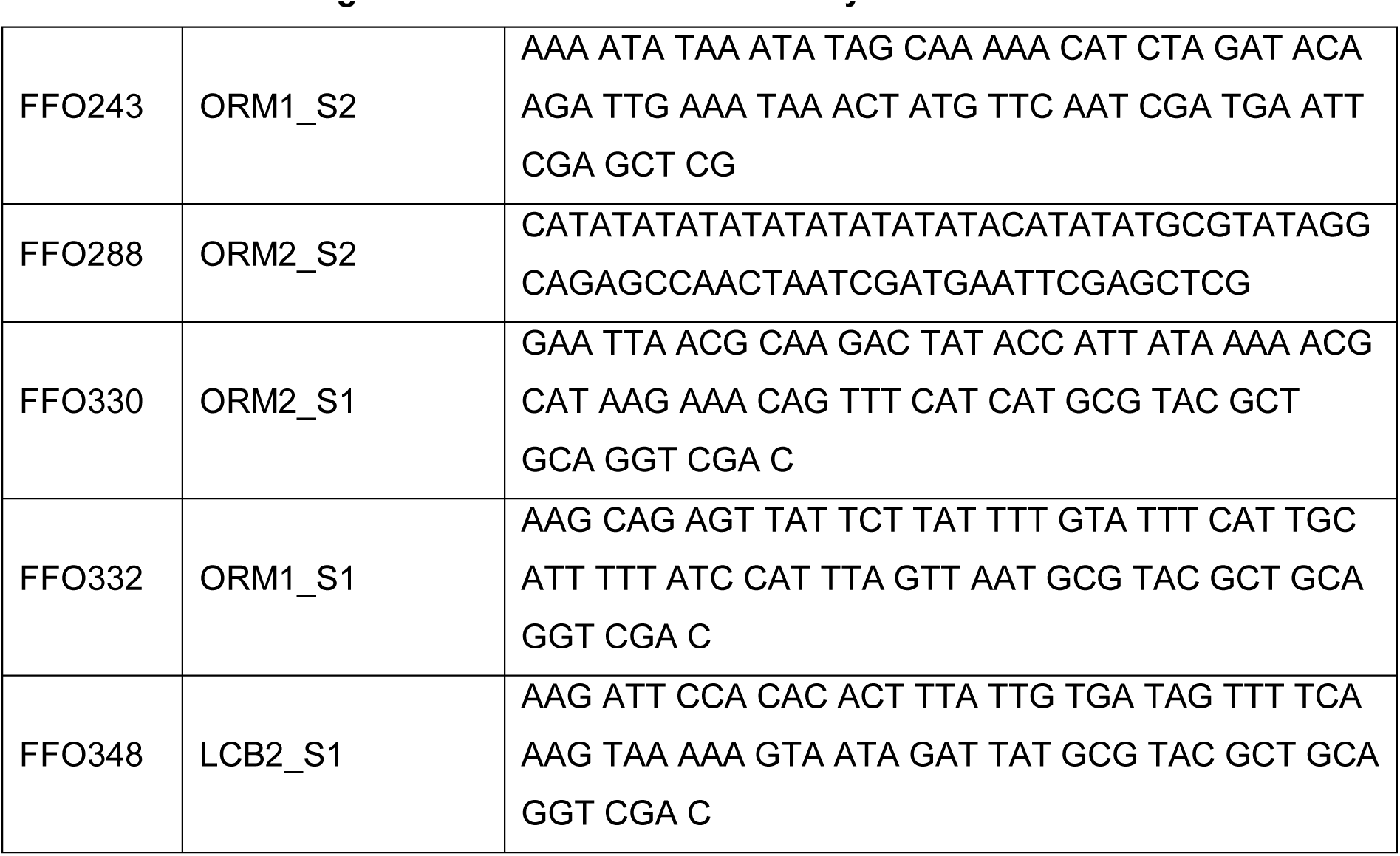
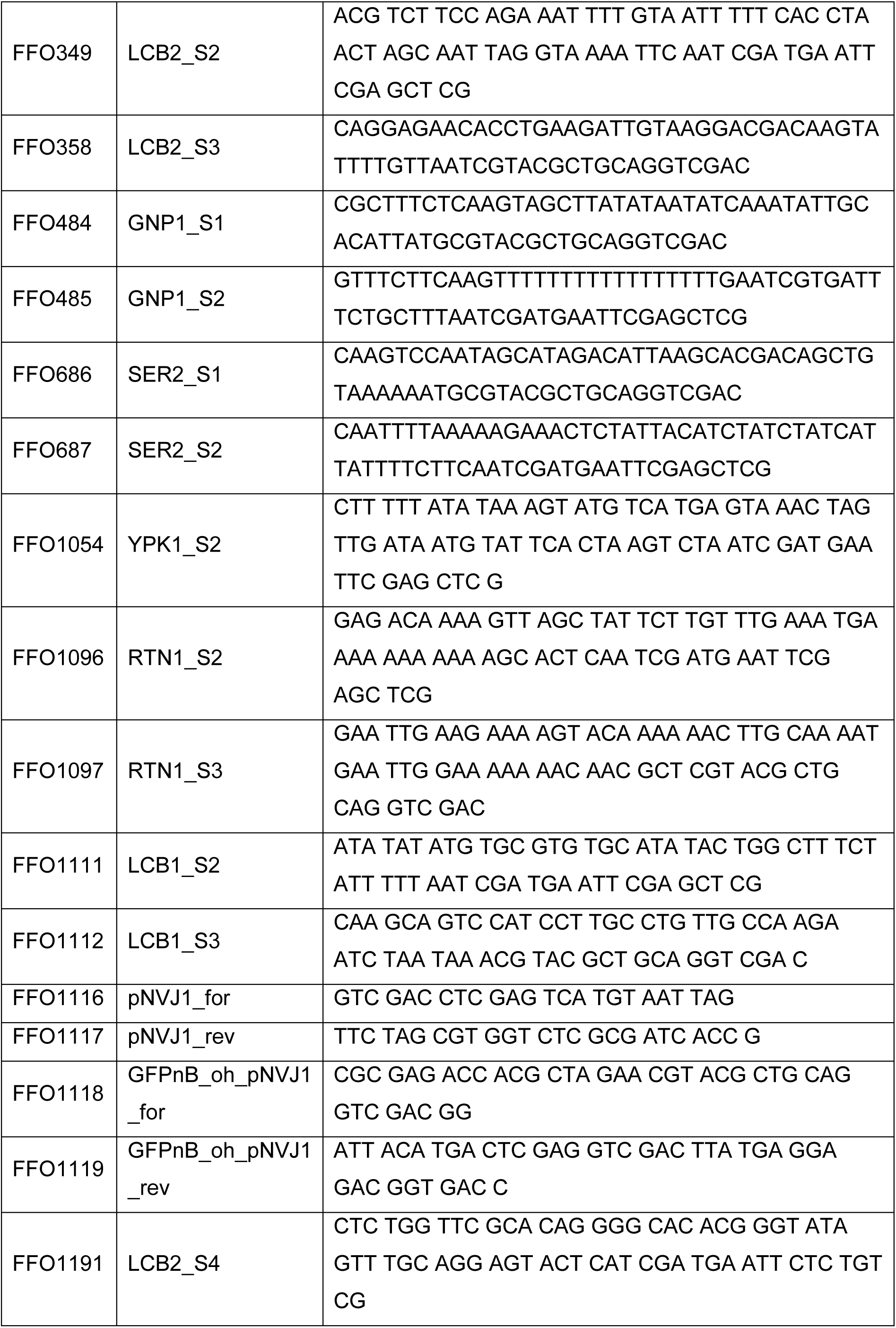
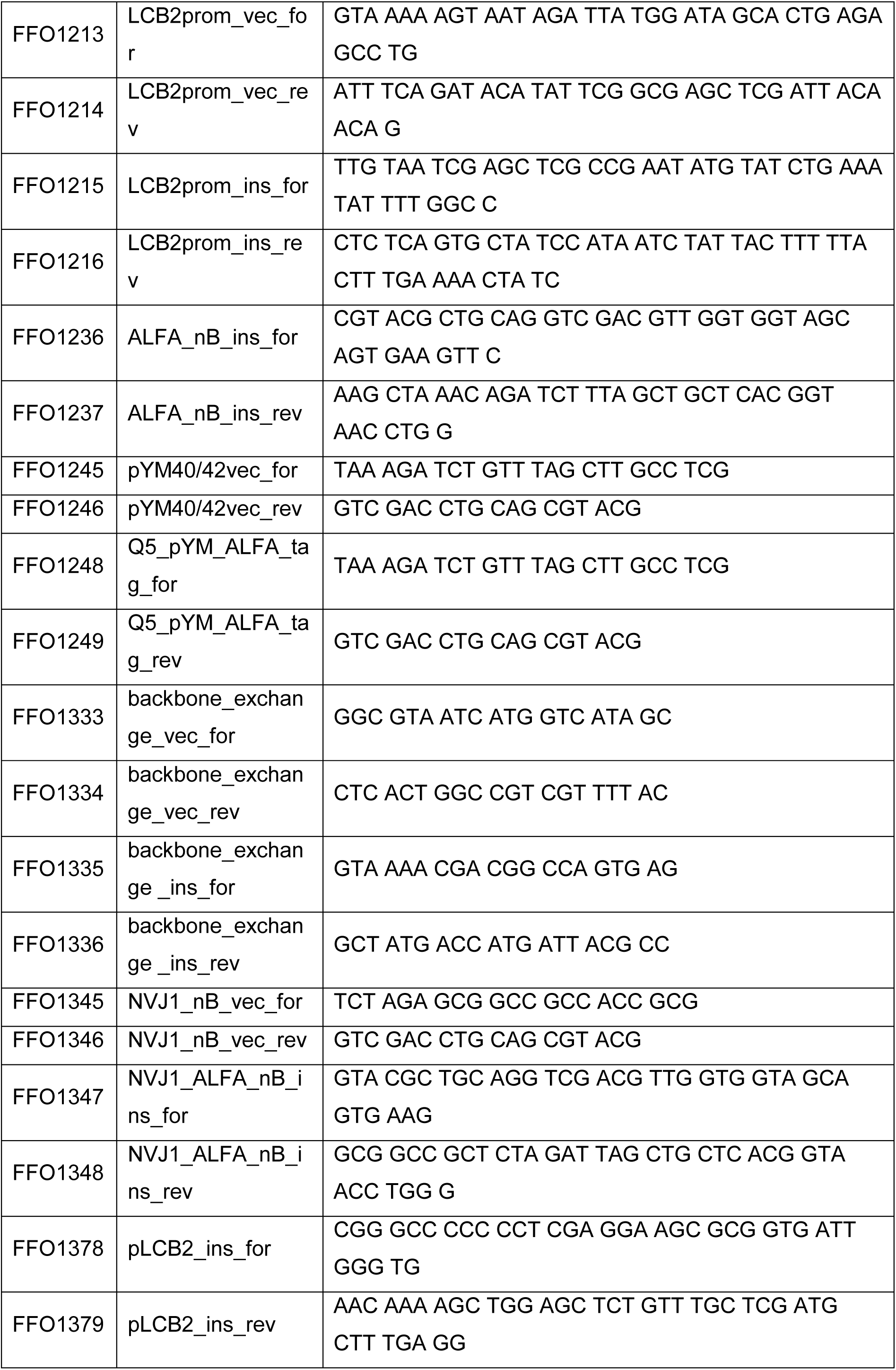
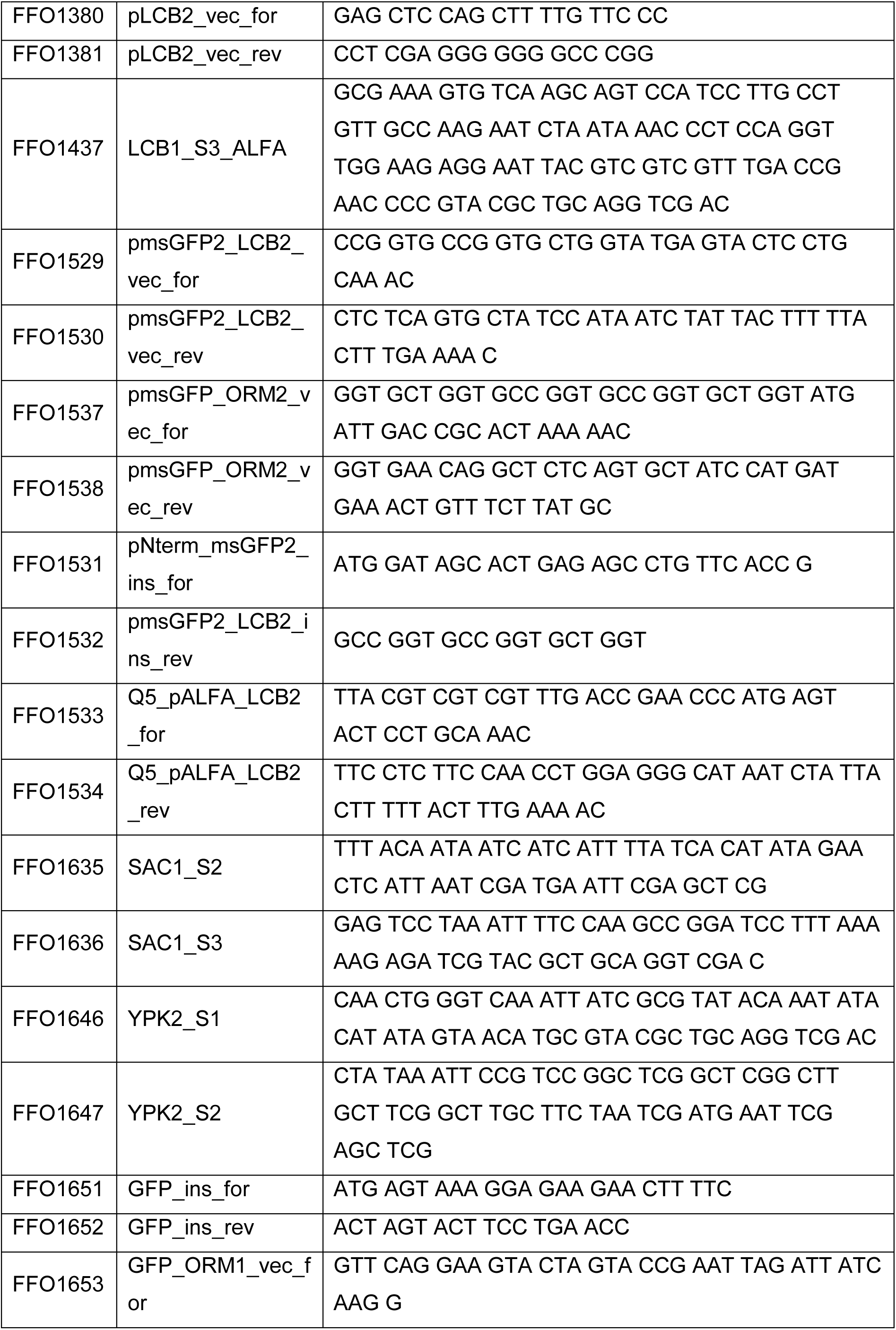
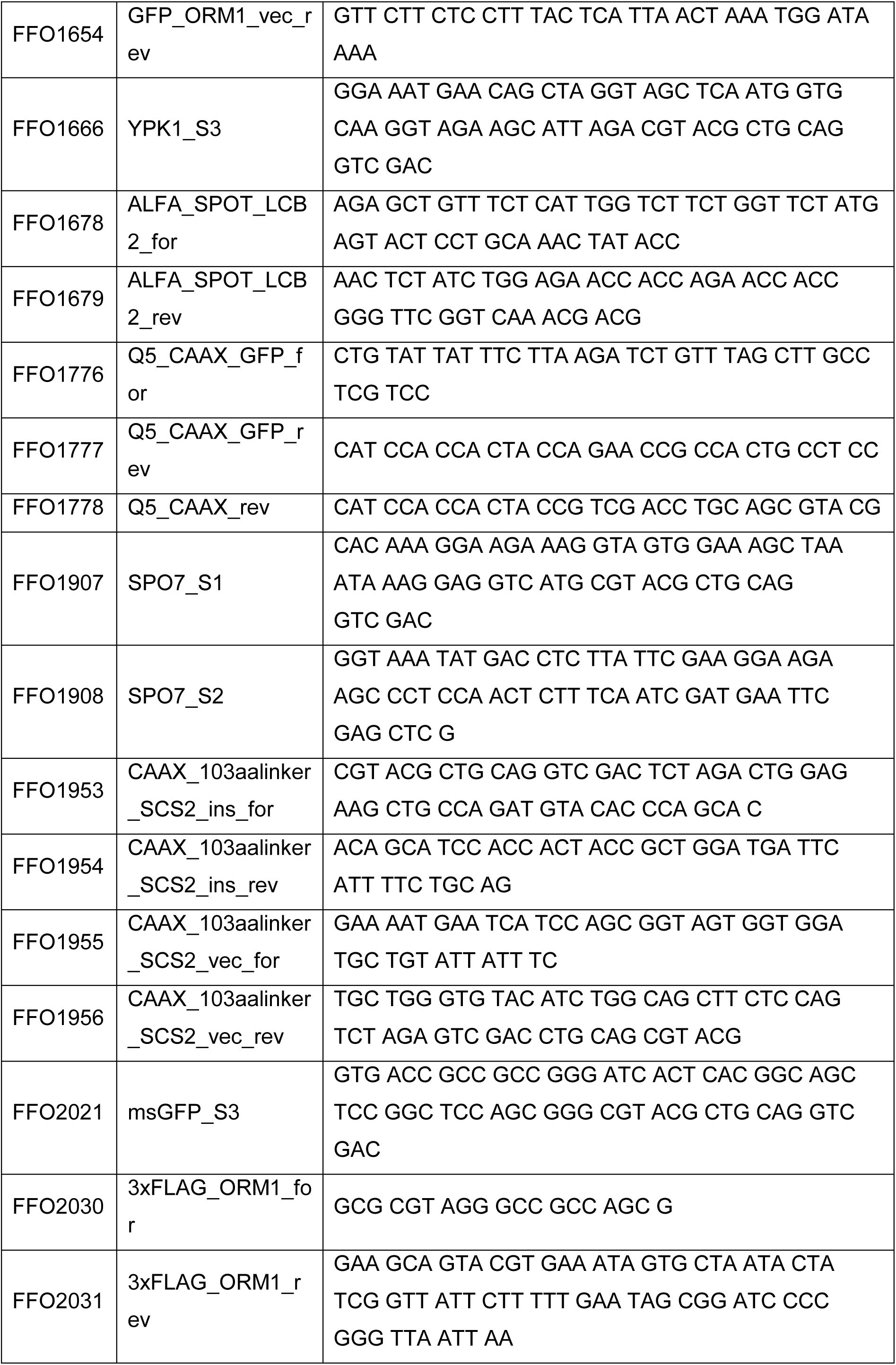
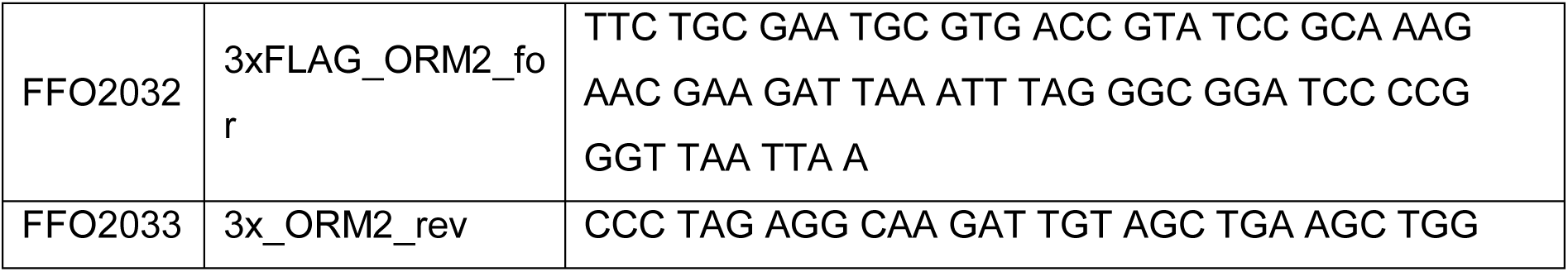
List of all oligonucleotides used in this study.

### Genetic interactions

To conduct tetrad analyses, diploid yeast cells were collected by centrifugation and placed onto 1% potassium acetate agar for sporulation at 30°C. After 3-5 days and microscopic inspection for ascus formation, a sample of each culture was suspended in 100 µL of sterile water. 5 µl of Zymolyase 20T (10 mg/mL; MP Biomedicals, Eschwege, Germany) were added, and incubated at room temperature for 9 minutes. A small amount of cells was streaked out on YPD plates and spores were segregated using a Singer MSM400 micromanipulator (Singer Instruments, Somerset, UK). The plates were then incubated for 3 days at 30°C.

### Spotting assays

For spotting assays cells from an overnight preculture were inoculated and grown to exponential growth phase in YPD. They were serial diluted and spotted to the YPD plates with and without addition of the indicated concentrations of myriocin (Sigma Aldrich). Plates were incubated for 2 days at 30 °C.

### Fluorescence microscopy

For fluorescence microscopy experiments cells were inoculated from an overnight preculture and grown to exponential growth phase. For the experiment with diploid cells (Fig. 3b) cells were imaged with equipment as described in (Eising *et al*, 2022). All other microscopy experiments were performed using an Axioscope 5 FL (Zeiss) microscope. It was equipped with an Axiocam 702 mono camera Plan-Apochromat 100x (1.4 numerical aperture (NA)) and an oil immersion objective using the ZEN 3.1 pro software. ImageJ was used for picture processing. Pictures were taken at same settings and were processed the same if not mentioned otherwise.

### Proteomics analysis

For proteomic analysis cells were inoculated from an overnight preculture and grown in YPD at 30 ° C until they reached exponential growth phase in triplicates. 2 OD units of all cultures were taken and pelleted at 4000 rpm for 2 minutes. Cell pellets were further treated according to the “iST Sample Preparation Kit (Pelleted cells & precipitated protein)” protocol with the iST Sample Preparation Kit (Preomics) for cells lysis and protein digestion. Dried peptides were resuspended in 50 µl LC-Load and 3 µl were loaded for LC-MS/MS measurement on a Thermo Ultimate 3000 RSLC nano system connected to a Q ExactivePlus mass spectrometer (Thermo Fisher Scientific) as described before (Limar *et al*, 2023). Briefly, resulting peptides were transferred to a glass vial and 3 µl were used to perform reversed-phase chromatography on a Thermo Ultimate 3000 RSLC nano system connected to a QExactivePLUS mass spectrometer (Thermo Fisher Scientific) through a nano-electrospray ion source. Peptides were separated on a PepMap RSLC C18 easy spray column (2 µm, 100 Å, 75 µm x 50 cm, Thermo Fisher Scientific) with an inner diameter of 75 µm. The column temperature was kept at 40 °C. The peptides were eluted from the column via a linear gradient of acetonitrile from 12-35% in 0.1% formic acid for 80 min at a constant flow rate of 200 nl/min followed by a 20 min increase to 60% and finally 10 min to reach 90% buffer B. Eluted peptides from the column were directly electro sprayed into the mass spectrometer. Mass spectra were acquired on the Q ExactivePlus in a data-dependent mode to automatically switch between full scan MS and up to ten data-dependent MS/MS scans. The maximum injection time for full scans was 50 ms, with a target value of 3,000,000 at a resolution of 70,000 at m/z 200. The ten most intense multiply charged ions (z≥2) from the survey scan were selected with an isolation width of 1.6 Th and fragment with higher energy collision dissociation with normalized collision energies of 27. Target values for MS/MS were set at 100,000 with a maximum injection time of 80 ms at a resolution of 17,500 at m/z 200. To avoid repetitive sequencing, the dynamic exclusion of sequenced peptides was set at 20 s. Resulting data were analyzed with MaxQuant (V2.1.4.0, www.maxquant.org) (Cox & Mann, 2008; Cox *et al*, 2011) and Perseus (V2.0.7.0, www.maxquant.org/perseus) (Tyanova *et al*, 2016).

### Pulldown experiments

For pull down experiments, cells were inoculated from an overnight preculture in 100 ml YPD in triplicates and grown to exponential growth phase at 30 °C. Same amounts of cells were harvested from all cultures at 4000 rpm 4°C for 5 min and snap frozen as cell pellets in Eppendorf tubes. Cells were lysed in with glass beads in 500 µl SPOT PD buffer (20mM HEPES pH 7.4, 150mM KOAc, 5% Glycerol, 1% GDN, Roche Complete Protease Inhibitor Cocktail EDTA free, Roche) using a FastPrep (MP biomedicals). Supernatant was cleared at 14000 rpm for 10 minutes and incubated for 30 minutes rotating at 4 °C together with 25 µl pre-equilibrated Spot-Trap beads (Chromotek). Beads were washed 4 times with GFP PD buffer at 2500 g for 2 minutes at 4 ° C. Afterwards, they were washed two times with Wash buffer (20mM HEPES pH 7.4, 150mM KOAc, 5% Glycerol) at 2500 g for 2 minutes at 4 °C. Beads were further treated following the “iST Sample Preparation Kit (Agarose Immunoprecipitation Samples)” protocol with the iST Sample Preparation Kit (Preomics) for protein digestion. Dried peptides were resuspended in 10 µl LC-Load and 5 µl were loaded for LC-MS/MS analyzes using the same settings and evaluation methods as described above. Resulting data were analyzed with MaxQuant (V2.0.3.0, www.maxquant.org) (Cox & Mann, 2008; Cox *et al*, 2011) and Perseus (V2.0.7.0, www.maxquant.org/perseus) (Tyanova *et al*, 2016).

### Lipidomics

Cultures were inoculated from a logarithmic growing preculture in 50 ml YPD and grown to exponential growth phase at 30 °C. Cells were collected at 4000 rpm 5 min 4 °C and snap frozen in liquid nitrogen. Cells were lysed with glass beads in 500 µl 155 mM ammonium formiate using a FastPrep (MP biomedicals). Lipid were extracted corresponding to 400 µg protein by two-step extraction as described previously (Ejsing *et al*, 2009). Internal standard (PG 17:0-14:1, PS 17:0-14:1, PE 17:0-20:4, PC 17:0-20:4, PI 17:0-20:4, LPE 17:1; Avanti Polar Lipids) was added before lipid extraction. First, lipids were extracted using 15:1 chloroform/methanol, which were later analysed via LC-MS/MS using positive ion mode. The remaining hydrophilic phase was re-extracted using 2:1 chloroform/methanol, which were later analysed via LC-MS/MS using negative ion mode. Dried lipids were dissolved in 50 µl 65:35 Buffer A (50:50 acetonitril/H_2_0, 10 mM ammonium formiate and 0.1% formic acid)/Buffer B (88:10:2 2-propanol/acetonitrile/H_2_0, 2 mM ammonium formiate and 0.02% formic acid). A C18 reverse-phase column (Thermo Accucore RP-MS, C18, 2.1 x 150 mm, 2.6 µm; Thermo Fisher Scientific) was used with a Shimadzu Nexera HPLC system with a heated electrospray ionization (HESI) and a ExactiveP*lus* Orbitrap mass spectrometer as described previously (Esch *et al*, 2020). The elution was performed with a 20-minute gradient. At 0 to 1 min, elution starts with 30% B and increases to 100% over 12 mins in a linear gradient. For 3 minutes 100 % B is maintained. Afterwards, solvent B was decreased to 30%. For 4 minutes 30% B is maintained for column re-equilibration. Flowrate was set to 0.3 ml/min. MS spectra of lipids were acquired in full-scan/data-dependent MS2 mode. The maximum injection time for full scans was 100 ms, with a target value of 3 000 000 at a resolution of 70 000 at m/z 200 with a mass range of 200–2000 m/z in both, positive and negative ion mode. The 10 most intense ions from the survey scan were selected and fragmented with HCD with a normalized collision energy of 27. Target values for MS/MS were set at 100 000 with a maximum injection time of 50 ms at a resolution of 17 500 at m/z 200. To avoid repetitive sequencing, the dynamic exclusion of sequenced lipids was set at 10 s. Resulting spectra were analyzed using LipidSearch 5.0 (Thermo Fischer). Lipid species were identified by database (>1,500,000 entries) search of positive (+H^+^; +NH_4_^+^) or negative (-H^-^; +HCOO^-^) adducts. Sample alignment was conducted with a retention time window of 0.5 minutes. Lipid standards were used for the calculation of lipid concentrations (PG 17:0-14:1, PS 17:0-14:1, PE 17:0-20:4, PC 17:0-20:4, PI 17:0-20:4, PA 15:0-18:1, LPS 17:1, LPE 17:1, LPC 17:1, LPI 17:1, TAG 15:0-18:1-15:0, DG 15:0-18:1 (Avanti Polar Lipids)). A LPE (negative) or Cer (positive) standard was used for the normalization between samples. Values are depicted as fold change from control strain.

### Flux analysis

Flux analysis was adapted from (Esch *et al*, 2020; Martínez-Montañés *et al*, 2020). Cultures were inoculated from a logarithmic growing preculture in 20 ml YPD and grown to exponential growth phase at 30 °C until they reach an OD_600_ of 0.8. Into 10 ml of the cultures [^13^C_3_^15^N_1_]-serine (CCN3000P1; CortecNet) to a final concentration of 3.8 mM was added (t=0). Samples (2.5 OD units each) were collected after 5, 15 and 30 minutes (t=5, 15, 30) at 4000 rpm 2 min 4 °C and cell pellets were directly snap frozen in liquid nitrogen. Lipids were extracted and analyzed as described in the targeted LCB and ceramide analysis paragraph.

### Heat shock experiments for LCB and ceramide analysis

Cultures were inoculated from a logarithmic growing preculture in 15 ml YPD and grown to exponential growth phase at 23 °C until they reach an OD_600_ of 0.8. Cells were splitted. 2.5 OD units of the culture were incubated at 23 °C for 5 minutes (no heat shock), another time 2.5 OD units of the culture were incubated at 39 °C for 5 minutes (heat shock) in a water bath. If mentioned, [^13^C_3_^15^N_1_]-serine (CCN3000P1; CortecNet) to a final concentration of 3.8 mM was added (t=0) before incubation. After heat shock, cells were harvested at 4000 rpm for 2 minutes, supernatant was poured out and cells were directly snap frozen in liquid nitrogen. Lipids were extracted and analyzed as described in the targeted LCB and ceramide analysis paragraph.

### LCB and ceramide analysis

Cells were thawed on ice and washed with ice-cold 155 mM ammonium formiate. Cell pellets were spiked with internal standard (Sphingosine d17:1, Ceramide d17:1/24:0; Avanti Polar Lipids) and lipid extraction with 2:1 chloroform/methanol was performed as described previously (Ejsing *et al*, 2009; Esch *et al*, 2020). Dried lipids were dissolved in 65:35 Buffer A (50:50 acetonitril/H_2_0, 10 mM ammonium formiate and 0.1% formic acid)/Buffer B (88:10:2 2-propanol/acetonitrile/H_2_0, 2 mM ammonium formiate and 0.02% formic acid). An external standard curve was prepared using dihydrosphingosine (DHS; Avanti Polar Lipids) 18:0, phytosphingosine (PHS; Avanti Polar Lipids) 18:0 and Ceramide t18:0/24:0 (Avanti Polar Lipids/Cayman). Samples were analyzed on a QTRAP 5500 LC-MS/MS (SCIEX) mass spectrometer connected to a Shimadzu Nexera HPLC system and an Accucore C30 LC column (150 mm x 2.1 mm 2.6 µm Solid Core; Thermo Fisher Scientific) in positive mode. For the gradient 40% B for 0.1 min was used. Followed by its increase from 40% to 50% over 1.4 min. Afterwards, Buffer B was increased from 50% to 100% over 1.5 min. 100% B was kept for 1 min and decreased to 40 % B for 0.1 min. 40% B was kept until the end of the gradient. A constant flow rate of 0.4 ml/min was used with a total analysis time of 6 minutes and an injection volume of 2 μl. The MS data were measured in positive ion mode, scheduled MRM mode without detection windows (Table4). For evaluation the SciexOS software was used. Internal standard was used for normalization. Measured OD_600_ units were used for correction of the used cell number.

**Table 4:** List of used transitions for targeted lipidomics.

| Description | Q1<br>mass | Q2<br>mass | CE (V) | CXP (V) | DP (V) | EP (V) |
| --- | --- | --- | --- | --- | --- | --- |
| Ceramide d17:1/24:0 | 636.602 | 249.8 | 47 | 13 | 120 | 10 |
| Phytoceramide 44:0;4 | 712.682 | 282.4 | 50 | 5 | 100 | 5 |
| *Phytoceramide 44:0;4 | 715.682 | 285.4 | 50 | 5 | 100 | 5 |
| Sphingosine d17:1 | 286.523 | 69 | 55 | 10 | 51 | 10 |
| Phytosphingosine 18:0 | 318.290 | 60 | 45 | 10 | 166 | 10 |
| Phytosphingosine 20:0 | 346.320 | 60 | 45 | 10 | 166 | 10 |
| *Phytosphingosine 18:0 | 321.290 | 63 | 45 | 10 | 166 | 10 |
| *Phytosphingosine 20:0 | 349.320 | 63 | 45 | 10 | 166 | 10 |
| Dihydrosphingosine 18:0 | 302.540 | 60 | 23 | 8 | 66 | 10 |
| Dihydrosphingosine 20:0 | 330.329 | 60 | 23 | 8 | 66 | 10 |
| *Dihydrosphingosine 18:0 | 305.540 | 63 | 23 | 8 | 66 | 10 |
| *Dihydrosphingosine 20:0 | 333.329 | 63 | 23 | 8 | 66 | 10 |

### Western Blots

Cultures were inoculated from a preculture in 20 ml YPD and grown to exponential growth phase at 23 °C until they reach an OD_600_ of around 1. Cells were splitted into 10 ml each. On half was incubated at 23 °C for 5 minutes (no heat shock), another half of the culture were incubated at 39 °C for 5 minutes (heat shock) in a water bath. After heat shock, cells were harvested at 4000 rpm for 2 minutes, supernatant was poured out and cells were directly frozen in liquid nitrogen. Cells were lysed with glass beads in 250 µl RIPA buffer (25mM Tris/HCl pH 7.6, 150mM NaCl, 1% NP-40, 1% sodium deoxycholate, 0.1% SDS, Roche complete protease inhibitor cocktail; Roche PhosStop tablet) using a FastPrep (MP biomedicals). Supernatant was cleared at 4000 rpm for 5 minutes. Protein concentration was determined and similar amount of protein was heated at 60 °C for 5 minutes in Laemilie buffer with DTT. FLAG-tagged proteins were detected with a mouse anti-FLAG (Roche) antibody diluted 1:1000. Pgk1 was detected with a 1:20000 diluted mouse anti-Pgk1 (RRID:AB_2532235; Thermo Fischer) antibody. FLAG and Pgk1 antibodies were detected using a DyLight 800 coupled anti-mouse IgG secondary antibody (SA535521; Invitrogen). ALFA-tagged proteins were detected using an 1:1000 diluted rabbit anti-ALFA antibody (N1505, Nanotag) using a DyLight 800 coupled mouse anti-rabbit IgG secondary antibody (SA535571; Invitrogen).

### Statistical analysis

For statistical analysis a two-sided t-test was used: * < 0.05. For proteomic experiments statistics were done as described before (Cox and Mann., 2008; Cox et al., 2011; Tyanova et al., 2016).

## Acknowledgements

We thank members of the Fröhlich lab for valuable discussions. We also thank Jacob Piehler (Osnabrück) for providing the nanobody expression plasmids. This work was supported by the DFG (grant FR 3674/2-2) and the SFB1557. Florian Fröhlich is a member of the Heisenberg program (FR 3674/4-1). Oliver Schmidt is supported by the Austrian Science Fund (FWF) [P 36187-B].

## References

1. Berchtold D, Piccolis M, Chiaruttini N, Riezman I, Riezman H, Roux A, Walther TC & Loewith R (2012) Plasma membrane stress induces relocalization of Slm proteins and activation of TORC2 to promote sphingolipid synthesis. Nat Cell Biol 14: 542–547

2. Bhaduri S, Aguayo A, Ohno Y, Proietto M, Jung J, Wang I, Kandel R, Singh N, Ibrahim I, Fulzele A, et al (2023) An ERAD-independent role for rhomboid pseudoprotease Dfm1 in mediating sphingolipid homeostasis. The EMBO Journal 42: e112275

3. Breslow DK, Collins SR, Bodenmiller B, Aebersold R, Simons K, Shevchenko A, Ejsing CS & Weissman JS (2010) Orm family proteins mediate sphingolipid homeostasis. Nature 463: 1048–1053

4. Buede R, Rinker-Schaffer C, Pinto WJ, Lester RL & Dickson RC (1991) Cloning and characterization of LCB1, a Saccharomyces gene required for biosynthesis of the long-chain base component of sphingolipids. J Bacteriol 173: 4325–4332

5. Busto JV, Elting A, Haase D, Spira F, Kuhlman J, Schäfer-Herte M & Wedlich-Söldner R (2018) Lateral plasma membrane compartmentalization links protein function and turnover. EMBO J 37

6. Cartier A & Hla T (2019) Sphingosine 1-phosphate: Lipid signaling in pathology and therapy. Science 366: eaar 5551

7. Chen J, Zheng XF, Brown EJ & Schreiber SL (1995) Identification of an 11-kDa FKBP12-rapamycin-binding domain within the 289-kDa FKBP12-rapamycin-associated protein and characterization of a critical serine residue. Proc Natl Acad Sci USA 92: 4947–4951

8. Cox J & Mann M (2008) MaxQuant enables high peptide identification rates, individualized p.p.b.-range mass accuracies and proteome-wide protein quantification. Nature Biotechnology 26: 1367–1372

9. Cox J, Neuhauser N, Michalski A, Scheltema RA, Olsen J V. & Mann M (2011) Andromeda: A Peptide Search Engine Integrated into the MaxQuant Environment. Journal of Proteome Research 10: 1794–1805

10. Craene J-OD, Coleman J, de Martin PE, Pypaert M, Anderson S, Yates JR, Ferro-Novick S & Novick P (2006) Rtn1p Is Involved in Structuring the Cortical Endoplasmic Reticulum□D. Molecular Biology of the Cell 17

11. Davis DL, Gable K, Suemitsu J, Dunn TM & Wattenberg BW (2019) The ORMDL/Orm–serine palmitoyltransferase (SPT) complex is directly regulated by ceramide: Reconstitution of SPT regulation in isolated membranes. Journal of Biological Chemistry 294: 5146–5156

12. Davis DL, Mahawar U, Pope VS, Allegood J, Sato-Bigbee C & Wattenberg BW (2020) Dynamics of sphingolipids and the serine palmitoyltransferase complex in rat oligodendrocytes during myelination. Journal of Lipid Research 61: 505–522

13. Dickson RC, Nagiec EE, Skrzypek M, Tillman P, Wells GB & Lester RL (1997) Sphingolipids Are Potential Heat Stress Signals inSaccharomyces. Journal of Biological Chemistry 272: 30196–30200

14. D’mello NP, Childress AM, Franklin DS, Kale SP, Pinswasdi C & Jazwinski SM (1994) Cloning and characterization of LAG1, a longevity-assurance gene in yeast. Journal of Biological Chemistry 269: 15451–15459

15. Dreisewerd K, Bien T & Soltwisch J (2022) MALDI-2 and t-MALDI-2 Mass Spectrometry Imaging. In Mass Spectrometry Imaging of Small Molecules, Lee Y-J (ed) pp 21–40. New York, NY: Springer US

16. Eising S, Esch B, Wälte M, Vargas Duarte P, Walter S, Ungermann C, Bohnert M & Fröhlich F (2022) A lysosomal biogenesis map reveals the cargo spectrum of yeast vacuolar protein targeting pathways. Journal of Cell Biology 221

17. Ejsing CS, Sampaio JL, Surendranath V, Duchoslav E, Ekroos K, Klemm RW, Simons K & Shevchenko A (2009) Global analysis of the yeast lipidome by quantitative shotgun mass spectrometry. Proceedings of the National Academy of Sciences 106: 2136–2141

18. Esch BM, Limar S, Bogdanowski A, Gournas C, More T, Sundag C, Walter S, Heinisch JJ, Ejsing CS, André B, et al (2020) Uptake of exogenous serine is important to maintain sphingolipid homeostasis in Saccharomyces cerevisiae. PLOS Genetics 16: e1008745

19. Friedman JR, Kannan M, Toulmay A, Jan CH, Weissman JS, Prinz WA & Nunnari J (2018) Lipid Homeostasis Is Maintained by Dual Targeting of the Mitochondrial PE Biosynthesis Enzyme to the ER. Developmental Cell 44: 261–270.e6

20. Gable K, Slife H, Bacikova D, Monaghan E & Dunn TM (2000) Tsc3p Is an 80-Amino Acid Protein Associated with Serine Palmitoyltransferase and Required for Optimal Enzyme Activity. Journal of Biological Chemistry 275: 7597–7603

21. Gatta AT, Wong LH, Sere YY, Calderón-Noreña DM, Cockcroft S, Menon AK & Levine TP (2015) A new family of StART domain proteins at membrane contact sites has a role in ER-PM sterol transport. eLife 4: e07253

22. Götzke H, Kilisch M, Martínez-Carranza M, Sograte-Idrissi S, Rajavel A, Schlichthaerle T, Engels N, Jungmann R, Stenmark P, Opazo F, et al (2019) The ALFA-tag is a highly versatile tool for nanobody-based bioscience applications. Nat Commun 10: 4403

23. Gournas C, Gkionis S, Carquin M, Twyffels L, Tyteca D & André B (2018) Conformation-dependent partitioning of yeast nutrient transporters into starvation-protective membrane domains. Proc Natl Acad Sci USA 115

24. Guillas I (2001) C26-CoA-dependent ceramide synthesis of Saccharomyces cerevisiae is operated by Lag1p and Lac1p. The EMBO Journal 20: 2655–2665

25. Gulshan K, Shahi P & Moye-Rowley WS (2010) Compartment-specific Synthesis of Phosphatidylethanolamine Is Required for Normal Heavy Metal Resistance. MBoC 21: 443– 455

26. Han G, Gupta SD, Gable K, Bacikova D, Sengupta N, Somashekarappa N, Proia RL, Harmon JM & Dunn TM (2019) The ORMs interact with transmembrane domain 1 of Lcb1 and regulate serine palmitoyltransferase oligomerization, activity and localization. Biochimica et Biophysica Acta (BBA) - Molecular and Cell Biology of Lipids 1864: 245–259

27. Han S, Lone MA, Schneiter R & Chang A (2010) Orm1 and Orm2 are conserved endoplasmic reticulum membrane proteins regulating lipid homeostasis and protein quality control. Proc Natl Acad Sci USA 107: 5851–5856

28. Hornemann T, Wei Y & von Eckardstein A (2007) Is the mammalian serine palmitoyltransferase a high-molecular-mass complex? Biochemical Journal 405: 157–164

29. Ikeda A, Schlarmann P, Kurokawa K, Nakano A, Riezman H & Funato K (2020) Tricalbins Are Required for Non-vesicular Ceramide Transport at ER-Golgi Contacts and Modulate Lipid Droplet Biogenesis. iScience 23: 101603

30. Janke C, Magiera MM, Rathfelder N, Taxis C, Reber S, Maekawa H, Moreno-Borchart A, Doenges G, Schwob E, Schiebel E, et al (2004) A versatile toolbox for PCR-based tagging of yeast genes: new fluorescent proteins, more markers and promoter substitution cassettes. Yeast 21: 947– 962

31. John Peter AT, Petrungaro C, Peter M & Kornmann B (2022a) METALIC reveals interorganelle lipid flux in live cells by enzymatic mass tagging. Nat Cell Biol 24: 996–1004

32. John Peter AT, Schie SNS, Cheung NJ, Michel AH, Peter M & Kornmann B (2022b) Rewiring phospholipid biosynthesis reveals resilience to membrane perturbations and uncovers regulators of lipid homeostasis. The EMBO Journal 41

33. Kajiwara K, Ikeda A, Aguilera-Romero A, Castillon GA, Kagiwada S, Hanada K, Riezman H, Muñiz M & Funato K (2013) Osh proteins regulate COPII-mediated vesicular transport of ceramide from the endoplasmic reticulum in budding yeast. Journal of Cell Science: jcs.132001

34. Klemm RW, Ejsing CS, Surma MA, Kaiser H-J, Gerl MJ, Sampaio JL, de Robillard Q, Ferguson C, Proszynski TJ, Shevchenko A, et al (2009) Segregation of sphingolipids and sterols during formation of secretory vesicles at the trans-Golgi network. Journal of Cell Biology 185: 601– 612

35. Klose C, Surma MA, Gerl MJ, Meyenhofer F, Shevchenko A & Simons K (2012) Flexibility of a Eukaryotic Lipidome – Insights from Yeast Lipidomics. PLoS ONE 7: e35063

36. Kvam E & Goldfarb DS (2006) Structure and function of nucleus-vacuole junctions: outer-nuclear-membrane targeting of Nvj1p and a role in tryptophan uptake. Journal of Cell Science 119: 3622–3633

37. Li C, Wen A, Shen B, Lu J, Huang Y & Chang Y (2011) FastCloning: a highly simplified, purification-free, sequence-and ligation-independent PCR cloning method. BMC Biotechnol 11: 92

38. Li S, Xie T, Liu P, Wang L & Gong X (2021) Structural insights into the assembly and substrate selectivity of human SPT–ORMDL3 complex. Nat Struct Mol Biol 28: 249–257

39. Limar S, Körner C, Martínez-Montañés F, Stancheva VG, Wolf VN, Walter S, Miller EA, Ejsing CS, Galassi VV & Fröhlich F (2023) Yeast Svf1 binds ceramides and contributes to sphingolipid metabolism at the ER cis-Golgi interface. J Cell Biol 222: e202109162

40. Liu L-K, Choudhary V, Toulmay A & Prinz WA (2017) An inducible ER–Golgi tether facilitates ceramide transport to alleviate lipotoxicity. Journal of Cell Biology 216: 131–147

41. Manford AG, Stefan CJ, Yuan HL, MacGurn JA & Emr SD (2012) ER-to-Plasma Membrane Tethering Proteins Regulate Cell Signaling and ER Morphology. Developmental Cell 23: 1129–1140

42. Martínez-Montañés F, Casanovas A, Sprenger RR, Topolska M, Marshall DL, Moreno-Torres M, Poad BLJ, Blanksby SJ, Hermansson M, Jensen ON, et al (2020) Phosphoproteomic Analysis across the Yeast Life Cycle Reveals Control of Fatty Acyl Chain Length by Phosphorylation of the Fatty Acid Synthase Complex. Cell Reports 32: 108024

43. van Meer G, Voelker DR & Feigenson GW (2008) Membrane lipids: where they are and how they behave. Nat Rev Mol Cell Biol 9: 112–124

44. Metterlein M, Yurlova L, Buchfellner A, Ruf B, Leingärtner V, Bogner J, Romer T, Kaiser PD, Hartlepp F & Linke-Winnebeck C (2018) Spot-Tag: a Nanobody-based Peptide-Tag System for Protein Detection, Purification and Imaging.

45. Millen JI, Pierson J, Kvam E, Olsen LJ & Goldfarb DS (2008) The Luminal N-Terminus of Yeast Nvj1 is an Inner Nuclear Membrane Anchor. Traffic 9: 1653–1664

46. Mondal K, Grambergs RC, Gangaraju R & Mandal N (2022) A Comprehensive Profiling of Cellular Sphingolipids in Mammalian Endothelial and Microglial Cells Cultured in Normal and High-Glucose Conditions. Cells 11: 3082

47. Muir A, Ramachandran S, Roelants FM, Timmons G & Thorner J (2014) TORC2-dependent protein kinase Ypk1 phosphorylates ceramide synthase to stimulate synthesis of complex sphingolipids. eLife 3: e03779

48. Nagiec MM, Baltisberger JA, Wells GB, Lester RL & Dickson RC (1994) The LCB2 gene of Saccharomyces and the related LCBJ gene encode subunits of serine palmitoyltransferase, the initial enzyme in sphingolipid synthesis. Proc Natl Acad Sci USA

49. Niles BJ & Powers T (2012) Plasma membrane proteins Slm1 and Slm2 mediate activation of the AGC kinase Ypk1 by TORC2 and sphingolipids in *S. cerevisiae*. Cell Cycle 11: 3745–3749

50. Olson DK, Fröhlich F, Christiano R, Hannibal-Bach HK, Ejsing CS & Walther TC (2015) Rom2-dependent Phosphorylation of Elo2 Controls the Abundance of Very Long-chain Fatty Acids. Journal of Biological Chemistry 290: 4238–4247

51. Robinson JB & Srere PA (1985) Organization of Krebs tricarboxylic acid cycle enzymes in mitochondria. Journal of Biological Chemistry 260: 10800–10805

52. Robinson JS, Klionsky DJ, Banta LM & Emr SD (1988) Protein sorting in Saccharomyces cerevisiae: isolation of mutants defective in the delivery and processing of multiple vacuolar hydrolases. Mol Cell Biol 8: 4936–4948

53. Roelants FM, Breslow DK, Muir A, Weissman JS & Thorner J (2011) Protein kinase Ypk1 phosphorylates regulatory proteins Orm1 and Orm2 to control sphingolipid homeostasis in *Saccharomyces cerevisiae*. Proc Natl Acad Sci USA 108: 19222–19227

54. Schäfer J-H, Körner C, Esch BM, Limar S, Parey K, Walter S, Januliene D, Moeller A & Fröhlich F (2023) Structure of the ceramide-bound SPOTS complex Biochemistry

55. Schmidt O, Weyer Y, Baumann V, Widerin MA, Eising S, Angelova M, Schleiffer A, Kremser L, Lindner H, Peter M, et al (2019) Endosome and Golgi-associated degradation (EGAD) of membrane proteins regulates sphingolipid metabolism. EMBO J 38

56. Shimobayashi M, Oppliger W, Moes S, Jenö P & Hall MN (2013) TORC1-regulated protein kinase Npr1 phosphorylates Orm to stimulate complex sphingolipid synthesis. MBoC 24: 870–881

57. Siniossoglou S (1998) A novel complex of membrane proteins required for formation of a spherical nucleus. The EMBO Journal 17: 6449–6464

58. Soltwisch J, Kettling H, Vens-Cappell S, Wiegelmann M, Müthing J & Dreisewerd K (2015) Mass spectrometry imaging with laser-induced postionization. Science 348: 211–215

59. Sotolongo Bellón J, Birkholz O, Richter CP, Eull F, Kenneweg H, Wilmes S, Rothbauer U, You C, Walter MR, Kurre R, et al (2022) Four-color single-molecule imaging with engineered tags resolves the molecular architecture of signaling complexes in the plasma membrane. Cell Reports Methods 2: 100165

60. Stradalova V, Blazikova M, Grossmann G, Opekarová M, Tanner W & Malinsky J (2012) Distribution of Cortical Endoplasmic Reticulum Determines Positioning of Endocytic Events in Yeast Plasma Membrane. PLoS ONE 7: e35132

61. Su W-M, Han G-S, Dey P & Carman GM (2018) Protein kinase A phosphorylates the Nem1–Spo7 protein phosphatase complex that regulates the phosphorylation state of the phosphatidate phosphatase Pah1 in yeast. Journal of Biological Chemistry 293: 15801–15814

62. Sun Y, Miao Y, Yamane Y, Zhang C, Shokat KM, Takematsu H, Kozutsumi Y & Drubin DG (2012) Orm protein phosphoregulation mediates transient sphingolipid biosynthesis response to heat stress via the Pkh-Ypk and Cdc55-PP2A pathways. MBoC 23: 2388–2398

63. Tabuchi M, Audhya A, Parsons AB, Boone C & Emr SD (2006) The Phosphatidylinositol 4,5-Biphosphate and TORC2 Binding Proteins Slm1 and Slm2 Function in Sphingolipid Regulation. Mol Cell Biol 26: 5861–5875

64. Tang X, Punch JJ & Lee W-L (2009) A CAAX motif can compensate for the PH domain of Num1 for cortical dynein attachment. Cell Cycle 8: 3182–3190

65. Traenkle B, Segan S, Fagbadebo FO, Kaiser PD & Rothbauer U (2020) A novel epitope tagging system to visualize and monitor antigens in live cells with chromobodies. Sci Rep 10: 14267

66. Tyanova S, Temu T, Sinitcyn P, Carlson A, Hein MY, Geiger T, Mann M & Cox J (2016) The Perseus computational platform for comprehensive analysis of (prote)omics data. Nature Methods 13: 731–740

67. Vallée B & Riezman H (2005) Lip1p: a novel subunit of acyl-CoA ceramide synthase. EMBO J 24: 730– 741

68. Voelker DR (1997) Phosphatidylserine decarboxylase. Biochimica et Biophysica Acta (BBA) - Lipids and Lipid Metabolism 1348: 236–244

69. Wadsworth JM, Clarke DJ, McMahon SA, Lowther JP, Beattie AE, Langridge-Smith PRR, Broughton HB, Dunn TM, Naismith JH & Campopiano DJ (2013) The Chemical Basis of Serine Palmitoyltransferase Inhibition by Myriocin. J Am Chem Soc 135: 14276–14285

70. Walther TC, Brickner JH, Aguilar PS, Bernales S, Pantoja C & Walter P (2006) Eisosomes mark static sites of endocytosis. Nature 439: 998–1003

71. Wang Y, Niu Y, Zhang Z, Gable K, Gupta SD, Somashekarappa N, Han G, Zhao H, Myasnikov AG, Kalathur RC, et al (2021) Structural insights into the regulation of human serine palmitoyltransferase complexes. Nat Struct Mol Biol 28: 240–248

72. West M, Zurek N, Hoenger A & Voeltz GK (2011) A 3D analysis of yeast ER structure reveals how ER domains are organized by membrane curvature. Journal of Cell Biology 193: 333–346

73. Zimmermann C, Santos A, Gable K, Epstein S, Gururaj C, Chymkowitch P, Pultz D, Rødkær SV, Clay L, Bjørås M, et al (2013) TORC1 Inhibits GSK3-Mediated Elo2 Phosphorylation to Regulate Very Long Chain Fatty Acid Synthesis and Autophagy. Cell Reports 5: 1036–1046

