## Supplementary Figures 1 and 2 for "Utilizing a nanobody recruitment approach for assessing serine palmitoyltransferase activity in ER sub-compartments of yeast"

### Supplementary material

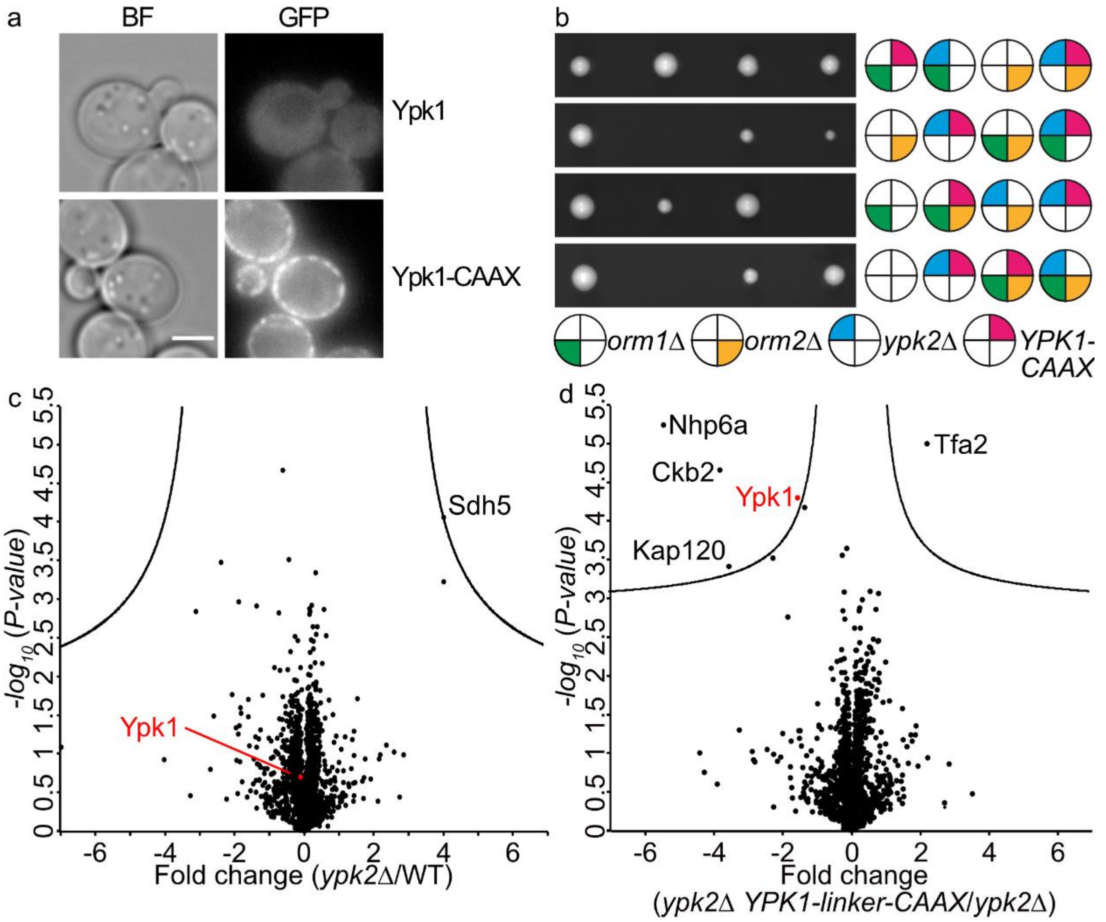


**Supplementary Figure 1: Evaluation of yeast cells with plasma membrane recruited Ypk kinases. (a)** Localization of wildtype GFP-tagged Ypk1 and (upper panel) and GFP-CAAX tagged Ypk1 (lower panel) are shown as representative mid-sections. Brightfield images (left panels) and fluorescent images are shown (right panels). Scale bar = 5 µm. **(b)** Plasma membrane targeted Ypk1 without linker is not functional. Tetrad analysis of *orm1Δ* (green) *orm2Δ* (yellow) *ypk2Δ* (blue) and *YPK1-CAAX* (pale red) cells. **(c,d)** Volcano plot identifying proteins that are enriched or de-enriched in Ypk1-linker CAAX cells. Fold changes were calculated from three independent experiments comparing the whole cell lysate proteome of *ypk2Δ* cells to WT cells **(c)** and *ypk2Δ YPK1-linker-CAAX* cells to *ypk2Δ* cells **(d)**. Fold changes were plotted on the x-axis against negative logarithmic P-values of the t-test performed from three replicates. Ypk1 is marked in red. The hyperbolic curve separates depleted proteins (left side) and enriched proteins (right side) from unaffected proteins (black dots).

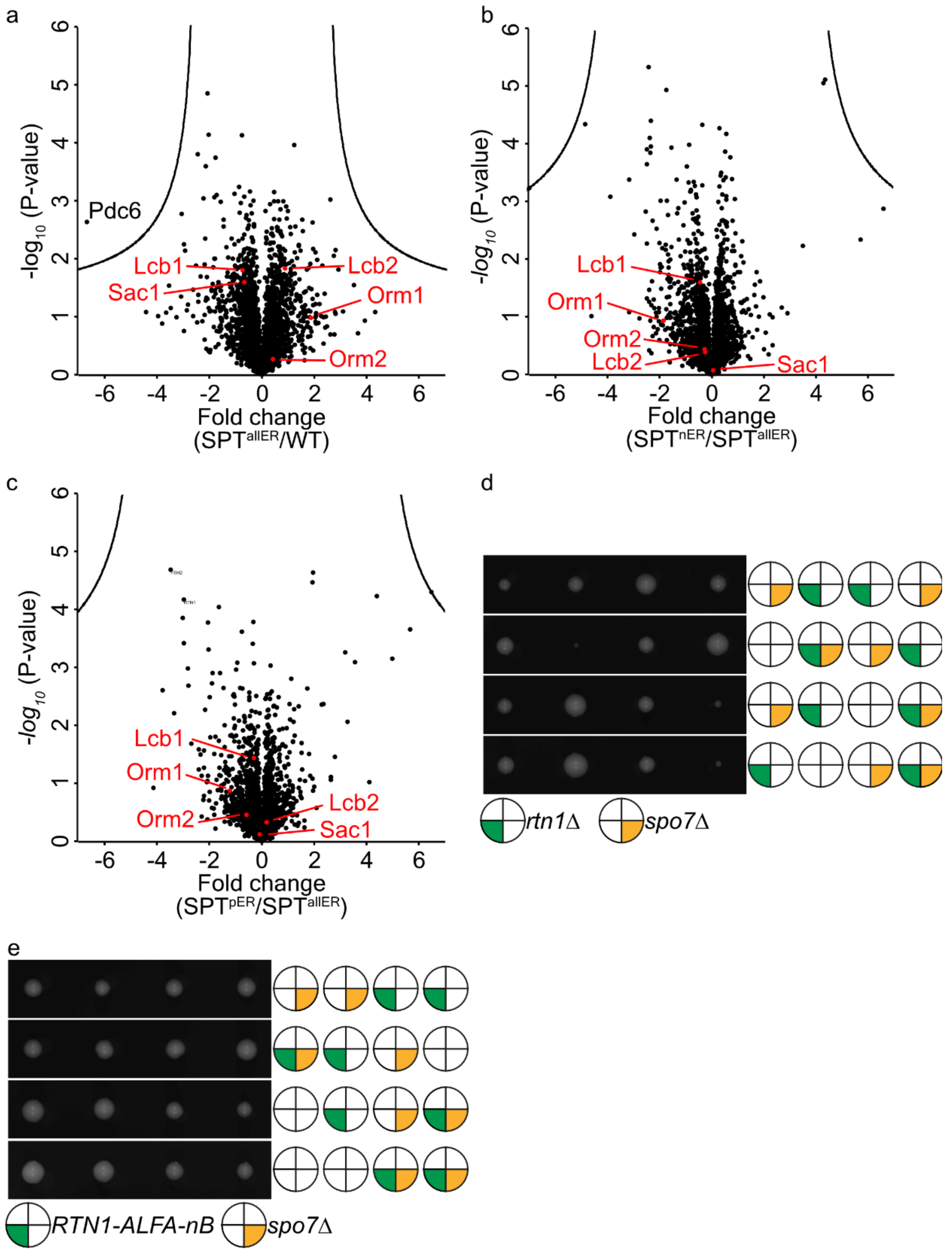


**Supplementary Figure 2: Evaluation of cellular effects after SPOTS recruitment.** Volcano plot identifying proteins that are enriched or de-enriched in SPT rewiring strains. Fold changes were calculated from three independent experiments comparing the whole cell lysate proteome of **(a)** $SPT^{allER}$ cells to WT cells, **(b)** $SPT^{nER}$ cells to $SPT^{allER}$ cells and **(c)** $SPT^{pER}$ cells to $SPT^{allER}$ cells. Fold changes were plotted on the x-axis against negative logarithmic P-values of the t-test performed from three replicates. SPOTS subunits are marked in red. The hyperbolic curve separates depleted proteins (left side) and enriched proteins (right side) from unaffected proteins (black dots). **(d)** Tetrad analysis of *rtn1Δ* (green) mutants crossed with *spo7Δ* mutants. **(e)** Tetrad analysis of *RTN1-ALFA-nB* (green) cells crossed with *spo7Δ* mutants.
